# VHL Mutation Drives Human Clear Cell Renal Cell Carcinoma Progression Through PI3K/AKT-Dependent Cholesteryl Ester Accumulation

**DOI:** 10.1101/2023.01.02.522447

**Authors:** Shuo Zhang, Tinghe Fang, Yexuan He, Weichen Feng, Zhuoyang Yu, Yaoyao Zheng, Chi Zhang, Shuai Hu, Zhuojun Liu, Jia Liu, Jian Yu, Han Zhang, Anbang He, Yanqing Gong, Zhisong He, Kaiwei Yang, Zhijun Xi, Wei Yu, Liqun Zhou, Lin Yao, Shuhua Yue

## Abstract

**Background:** Cholesteryl ester (CE) accumulation in intracellular lipid droplets (LDs) is an essential signature of clear cell renal cell carcinoma (ccRCC), but its molecular mechanism and pathological significance remain elusive.

**Methods:** Enabled by the label-free Raman spectromicroscopy, which integrated stimulated Raman scattering microscopy with confocal Raman spectroscopy on the same platform, we quantitatively analyzed LD distribution and composition at the single cell level in intact ccRCC cell and tissue specimens *in situ* without any processing or exogenous labeling. Since we found that commonly used ccRCC cell lines actually did not show the CE-rich signature, primary cancer cells were isolated from human tissues to retain the lipid signature of ccRCC with CE level as high as the original tissue, which offers a preferable cell model for the study of cholesterol metabolism in ccRCC. Moreover, we established a patient-derived xenograft (PDX) mouse model that retained the CE-rich phenotype of human ccRCC.

**Findings:** Surprisingly, our results revealed that CE accumulation was induced by tumor suppressor VHL mutation, the most common mutation of ccRCC. Moreover, VHL mutation was found to promote CE accumulation by upregulating HIFα and subsequent PI3K/AKT/mTOR/SREBPs pathway. Inspiringly, inhibition of cholesterol esterification remarkably suppressed ccRCC aggressiveness *in vitro* and *in vivo* with negligible toxicity, through the reduced membrane cholesterol-mediated downregulations of integrin and MAPK signaling pathways.

**Interpretation:** Collectively, our study improves current understanding of the role of CE accumulation in ccRCC and opens up new opportunities for treatment.

**Funding:** This work was supported by National Natural Science Foundation of China (No. U23B2046 and No. 62027824), National Key R&D Program of China (No. 2023YFC2415500), Fundamental Research Funds for the Central Universities (No. YWF-22-L-547), PKU-Baidu Fund (No. 2020BD033), Peking University First Hospital Scientific and Technological Achievement Transformation Incubation Guidance Fund (No.2022CX02), and Beijing Municipal Health Commission (No. 2020-2Z-40713).

## Introduction

As an essential hallmark of human cancers, dysregulated lipid metabolism promotes cancer development and progression through increased lipid biosynthesis, transport, oxidation, and hydrolysis ^1,2^. For accommodating vigorous lipid turnover, excessive lipids are esterified into neutral lipids, primarily triacylglycerols (TAGs) and cholesteryl esters (CEs), and subsequently stored in lipid droplets (LDs) ^3^. As a critical organelle to maintain intracellular lipid homeostasis, LD has been frequently found to accumulate in various cancers and play an important role in cancer development ^4,5^.

As the most common (>75%) and aggressive subtype of renal carcinoma, clear cell renal cell carcinoma (ccRCC) is known to be one of the most LD-rich cancers ^6,7^. A couple of studies have been conducted to elucidate the mechanism underlying the accumulation and function of LDs in ccRCC ^8–10^. Specifically, as the most prevalent molecular feature of ccRCC, constitutive stabilization of hypoxia inducible factors (HIFs) ^11^, induced by the mutation of tumor suppressor VHL (60∼90% of ccRCC) ^12^, was found to drive LD accumulation via downregulation of the rate-limiting enzyme of mitochondrial fatty acid transport, carnitine palmitoyltransferase 1A (CPT1A) ^8^, and upregulation of the LD-associated protein perilipin 2 (PLIN2) ^9^. Such LD accumulation was shown to maintain endoplasmic reticulum (ER) integrity ^9^ and promote lipid homeostasis by suppressing fatty acid toxicity ^10^, particularly under hypoxic stress, thus further facilitate ccRCC cell survival.

It is well known that the major form of neutral lipid accumulated in ccRCC is CE but not TAG ^7,13,14^. Unfortunately, most of the previous studies in ccRCC focus on fatty acid metabolism instead of cholesterol metabolism. More recently, high-density lipoprotein (HDL) has been found to be the source of cholesterol, the driving force for ccRCC growth ^15^. Nevertheless, the biological mechanism and pathological significance of CEs accumulated in ccRCC remain unclear.

The reason why CE accumulation in ccRCC has not been well studied is partially due to a lack of proper analytical tools. For LD visualization in single cells, lipophilic dyes are commonly used but lack the compositional information. For lipid compositional analysis, analytical methods, such as mass spectrometry and nuclear magnetic resonance spectroscopy, are regularly performed but cannot provide the spatial distribution information. Recently developed mass spectrometry imaging has been applied to chemical imaging of tissues *in situ* ^16^, but the limited spatial resolution prohibited its application in the study of LD biology. As a chemical imaging method with submicron resolution, stimulated Raman scattering (SRS) microscopy has made significant contributions to the study of lipid metabolism at the single cell level in various type of diseases ^17,18^, including cancer ^19–23^.

In this study, we elucidated the role of CE accumulation in ccRCC by integrating SRS-based spectroscopic imaging with biochemical and molecular biology methods. Our chemical imaging data revealed an aberrant accumulation of CEs in human ccRCC tissues (n = 24), but not in normal adjacent tissues (n = 24), collected from 41 patients undergone surgery. To unravel the underlying mechanism of CE accumulation, we firstly analyzed a panel of commonly used ccRCC cell lines, including 786-O, 769-P, OSRC, Caki-1, and RCC4, but unfortunately did not find any detectable CEs in LDs, suggesting that these cell lines may not be appropriate for the study of cholesterol metabolism. This finding partially explains why the role of CEs in ccRCC progression remains unclear. To develop an appropriate cell model, we then extracted primary cancer cells from human ccRCC tissues and verified that the CE levels of primary cells were close to their original ccRCC tissues. Notably, the CE level was significantly higher in those with VHL mutation than those without VHL mutation, suggesting the possible link between VHL mutation and CE accumulation. By integrating with biological methods, we unraveled that the aberrant CE accumulation in ccRCC was induced by the VHL mutation and subsequently upregulated PI3K/Akt/mTOR/SREBPs pathway in a HIFα-dependent manner. Importantly, we demonstrated that inhibition of cholesterol esterification effectively impaired ccRCC aggressiveness *in vitro* and in patient-derived xenograft (PDX) mouse model with negligible toxicity, likely due to the reduced membrane cholesterol mediated downregulations of the integrin and MAPK signaling pathways. These results collectively improve understanding of the role of CE accumulation in ccRCC and open up new opportunities for treatment.

## Methods

### Human Tissue Specimens

All tissues specimens were obtained from ccRCC patients undergone nephrectomy at Peking University First Hospital. This study was approved by the Institutional Review Boards of both Beihang University and Peking University First Hospital. Pathological examination was performed by experienced pathologists. Details were described in the Supplementary Experimental Procedures.

### Cell Cultures

All cell lines were obtained from the American Type Culture Collection. Primary cancer cells were isolated from fresh human ccRCC tissues according to the procedures in the previous literature ^24^. Details were described in the Supplementary Experimental Procedures.

### Label-free Raman Spectromicroscopy

Label-free Raman spectromicroscopy was performed on unstained frozen tissue slices (∼20 μm) and live cells. SRS images were obtained on a picosecond SRS microscope, with the laser-beating frequency tuned to the C-H stretching vibration band at 2,850 cm^-1^. Compositional analysis of individual LDs and autofluorescent granules was conducted by integration of SRS imaging and confocal Raman spectral analysis on a single platform. Two-photon fluorescence signals were acquired through a 520/40 nm bandpass filter for the images of autofluorescent granules in human tissues. Experiments of SRS imaging/Raman spectral acquisition and quantitative analysis of LD amount/composition were performed as described previously^25^. Details were described in the Supplementary Experimental Procedures.

#### Hyperspectral SRS microscopy and related quantitative analysis

The hyperspectral SRS imaging was performed as described previously ^26^. For quantification of LD amount, we employed spectral phasor analysis to perform LD detection according to the previous literatures^27,28^. We then used the ratiometric approach to quantify the CE percentage according to the literature^29^ Details were described in the Supplementary Experimental Procedures.

### Lipid Extraction and Measurements

Lipids in tissues were extracted and measured as described in the Supplementary Experimental Procedures.

### DNA Sequencing and RNA Sequencing

Genomic DNA was extracted and purified from primary cells according to manufacturer’s instruction of DNAzol™ reagent (Thermo Fisher Scientific, 10503027). RNA was extracted from primary cells according to manufacturer’s instruction of TRIzol™ reagent (Thermo Fisher Scientific, 15596026). Details were described in the Supplementary Experimental Procedure.

### RNA Interference, Immunoblotting, Immunofluorescence staining, Filipin and BODIPY staining

These experiments were performed as described in the Supplementary Experimental Procedure.

### Cell Viability Assay, Cell Cycle Analysis, Migration/Invasion Assay

Cell viability assay was performed with the MTT colorimetric assay (Thermo Fisher Scientific, M6494). Cell cycle analysis used PI staining (Thermo Fisher Scientific, P3566) and then was performed by flow cytometer (Peking University Health Science Center). Migration and invasion assays were performed in Transwell chambers coated with and without Matrigel (Corning, 354234). Details were described in the Supplementary Experimental Procedures.

### PDX mouse model

All animal care and experimental procedures were performed in accordance with the guidelines for care and use of laboratory animals. For *in vivo* anti-tumor efficacy, safety evaluation, pharmacokinetic, and biodistribution analyses, subcutaneous PDX mouse model of ccRCC was established as a previously described protocol ^30^. Details were described in the Supplementary Experimental Procedures.

### Statistical Analysis

One-way ANOVA and Student’s t test were used for comparisons between groups. p < 0.05 was considered statistically significant. All data was presented as mean ±standard error of the mean (SEM).

## Results

### Aberrant CE accumulation in human ccRCC tissues but not in normal adjacent tissues

By integrating SRS microscopy and confocal Raman spectroscopy, we performed *in situ* lipid imaging and analysis of 24 ccRCC tissues and 24 normal adjacent tissues, collected from 41 patients undergone kidney cancer surgery. By tuning the laser beating frequency to be resonant with the C-H stretching vibration at 2,850 cm^-1^, strong SRS signals arose from the lipid-rich structures, such as cell membranes and LDs, and in the meanwhile much weaker signals came from the lipid-poor structures, such as the nuclei. Such imaging contrast permit clear visualization of cellular morphology and intracellular LD accumulation in a label-free manner. As shown in Fig. 1a-d and S1a-d, the SRS images showed very similar morphological features compared to the hematoxylin and eosin (H&E) images of the adjacent tissue slices. It was found that the autofluorescent granules, known as the lipofuscins, were consistently present in all normal tissues but not in any of the ccRCC tissues (Fig. 1e, 1f, S1e, S1f, and Table S1). More importantly, abundant LD accumulation was observed in the ccRCC tissues but not in any of the normal tissues (Fig. 1e, 1f, S1e, S1f, and Table S1). Quantitatively, the LD amount calculated in the way of LD area fraction was 13.7% ±2.1% in the ccRCC tissues (Fig. 1g and Table S1).

**Fig. 1:**
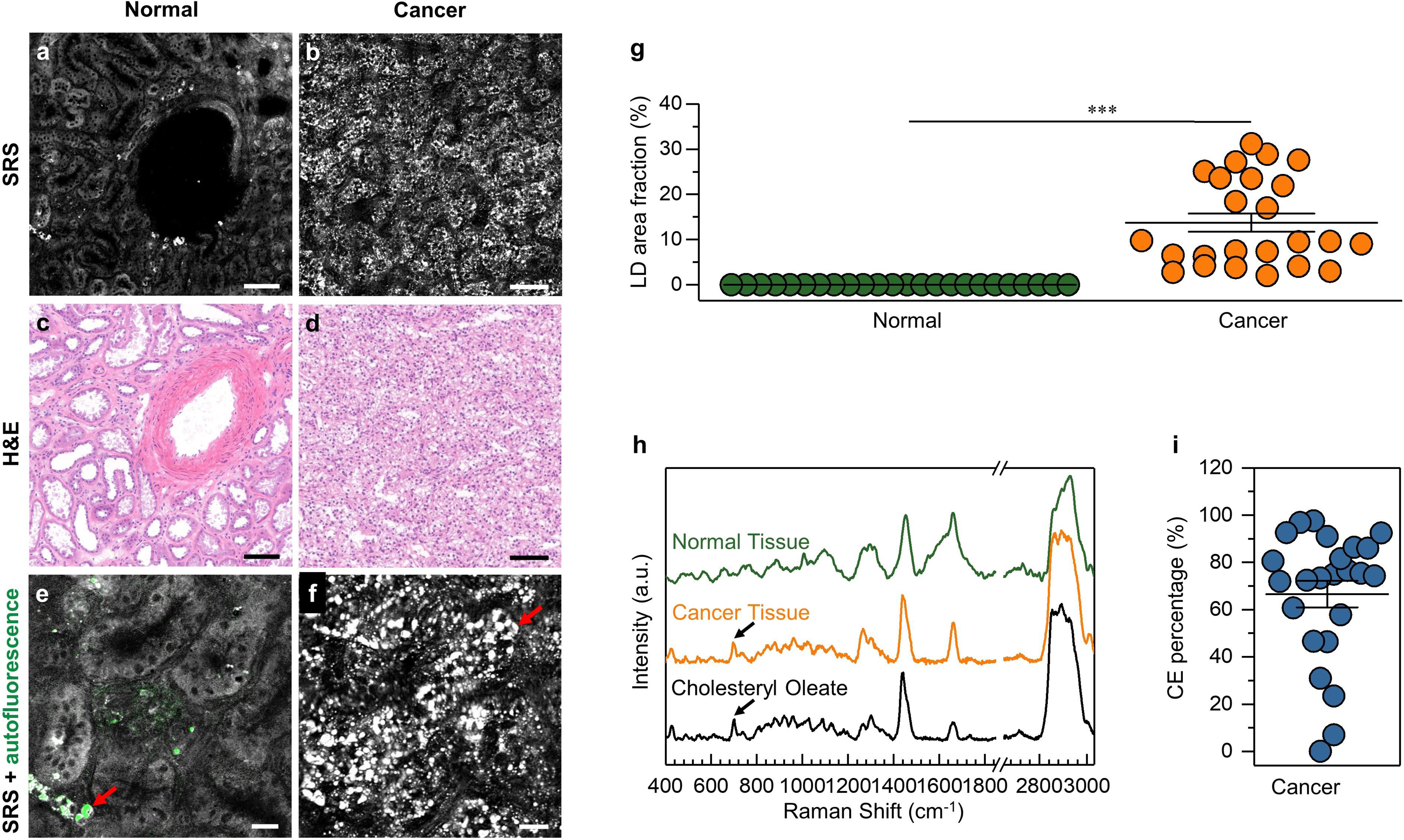
Aberrant CE accumulation in human ccRCC tissues. **(a, b)** Representative SRS images of human normal adjacent and ccRCC tissues. Scale bar, 100 µm. **(c, d)** H&E images of the adjacent slices shown in **a** and **b**. Scale bar, 100 µm. **(e, f)** Magnified SRS and autofluorescence images of the ones shown in **a** and **b**. Autofluorescent granules and LDs are indicated by red arrows. Scale bar, 25 µm. **(g)** Quantitation of LD area fraction for 24 normal adjacent and ccRCC tissues. Error bars represent SEM. ***p < 0.0005. **(h)** Representative Raman spectra of autofluorescence granule in normal adjacent tissue, LD in ccRCC tissue, and pure cholesteryl oleate. Spectral intensity was normalized by CH_2_ bending band at 1,442 cm^-1^. Black arrows indicate the band of cholesterol ring at 702 cm^-1^. **(i)** Quantitation of CE percentage for 24 ccRCC tissues. Error bars represent SEM.

For further compositional analysis by confocal Raman spectroscopy, the autofluorescent granules in normal tissues were found to show characteristic Raman peaks for lipid (1,200-1,800 cm^-1^), phenylalanine (1,000 cm^-1^), and prominent CH_3_ stretching (2,930 cm^-1^). In contrast, the LDs in ccRCC tissues exhibited a series of strong representative peaks for CEs, including cholesterol rings around 428 and 702 cm^-1^, ester bond around 1,742 cm^-1^ (Fig. 1h). Since LDs are primarily composed of TAGs and CEs, we made emulsions mixed with different molar ratios between cholesteryl oleate and glyceryl trioleate for Raman spectral analysis (Fig. S2a) to get a calibration curve for quantification of CE percentage in LDs ^23^. As shown in Fig. S2b, the height ratio between the most prominent band for cholesterol ring around 702 cm^-1^ and the CH_2_ bending band around 1,442 cm^-1^ was linearly proportional to the molar percentage of CEs in LDs. Based on this calibration curve, CE percentage in LDs of ccRCC tissues were as high as 66.5% ±5.7% (Fig. 1i and Table S1). The detailed quantifications of LD amount and composition for each patient are shown in the Table S1. Liquid chromatography-mass spectrometry analysis of lipids extracted from tissues confirmed that CE accumulation in ccRCC tissues was significantly higher than that in normal adjacent tissues (Fig. S3). These findings together demonstrate elevated CE accumulation and possible reprogramming of cholesterol metabolism in human ccRCC.

### Primary cell as a preferable model for the study of cholesterol metabolism in ccRCC

To investigate how CEs accumulate in human ccRCC, we employed SRS microscopy to quantitatively analyze lipid content in a panel of commonly used cell lines, including ccRCC cell lines (786-O, 769-P, Caki-1, OSRC, and RCC4), normal kidney epithelial HK-2 cell line and embryonic kidney 293 cell line. All the above ccRCC cell lines contained substantially fewer LDs than human cancer tissues and did not show significant differences compared to the normal kidney cell lines (Fig. 2a and 2b). Moreover, CEs were undetectable in any of the cell lines based on Raman spectral analysis (Fig. 2c). Our results suggest that these commonly used immortalized cell lines may not be suitable for the study of CE accumulation in ccRCC. In order to address this problem, primary cancer cells were isolated from human ccRCC tissues and confirmed by immunostaining of the epithelium marker (Fig. S4a). The SRS images revealed numerous LDs in primary cancer cells (Fig. 2a), also confirmed by Bodipy labeling (Fig. S4a), which represented the “clear cell” phenotype of ccRCC tissues. Quantitatively, LD amount in primary cancer cells was 10∼30 times greater than the cell lines (Fig. 2b). Based on Raman spectral analysis, the LDs in primary cancer cells showed CE levels (Fig. 2c) that were as high as the original cancerous tissues (Fig. S4b). In order to further confirm these findings, hyperspectral SRS imaging that could offer spectral information for each pixel on the image was established as decribed previously^26^. Based on spectral phasor analysis and ratiometric approach^26,29^, LD amount and CE percentage could be precisely visualized (Fig. S5a-S5d). As shown in Fig. S5e and S5f, the LD amount and CE level were found to be remarkably higher in primary cells compared to the cell lines, consistent with our previous measurements. Additionly, upon treatment of primary cell culture medium, LD amount was not changed in769-P, Caki-1, and RCC4, whereas significantly increased in 786-O and OSRC (Fig. S5g). Nevertheless, the CE level remained low without significant change in the tested ccRCC cell lines upon treatment of primary cell culture medium (Fig. S5h). These results confirm that culture medium is not the reason for the CE-rich phenotype of primary ccRCC cells. Collectively, our study indicates that primary cancer cell, but not cell line, is the appropriate cell model to elucidate cholesterol metabolic reprogramming in ccRCC.

**Fig. 2:**
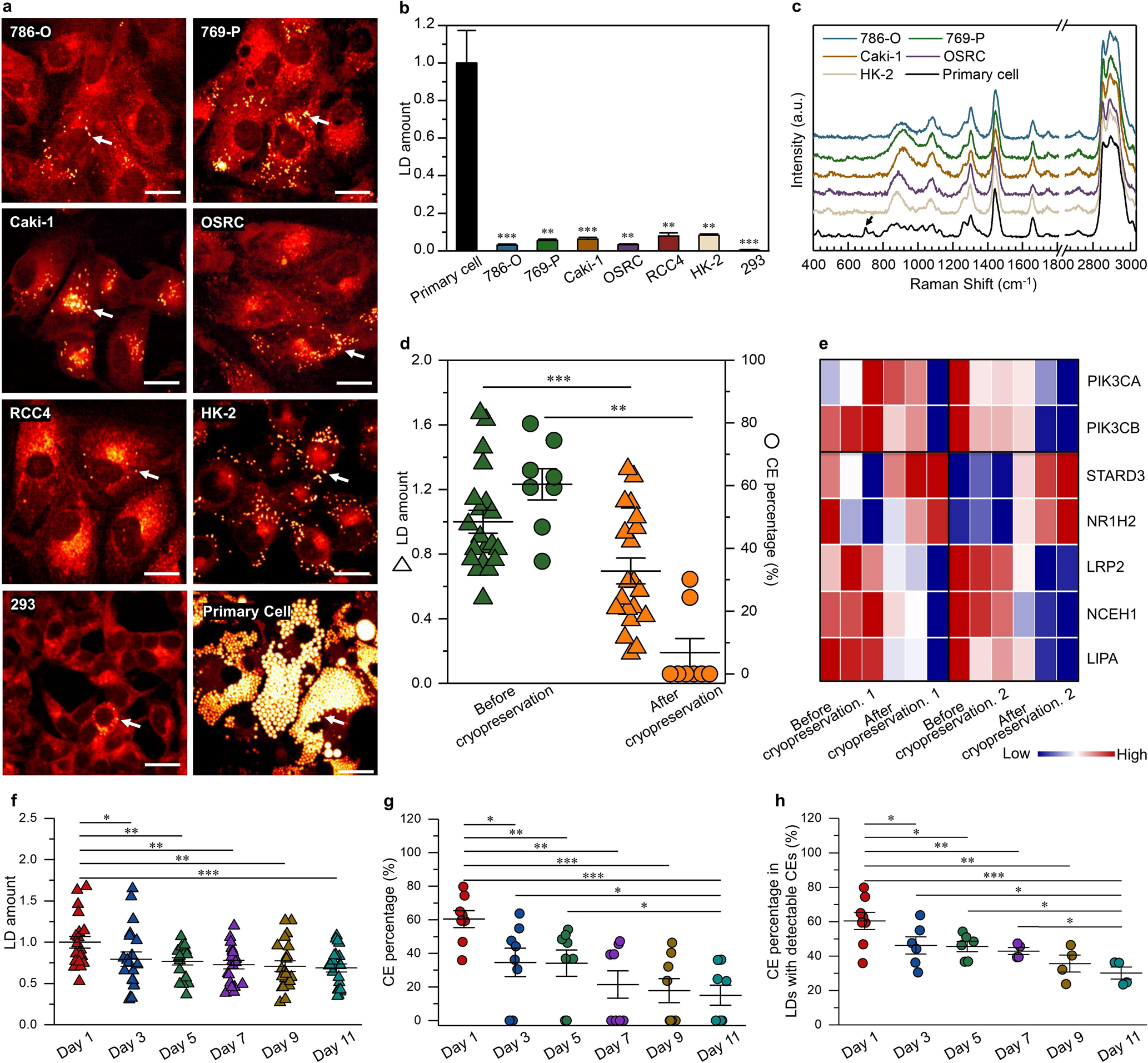
Primary cancer cell as an appropriate model for the study of cholesterol metabolism in ccRCC. **(a)** Representative SRS images of ccRCC cell lines (786-O, 769-P, Caki-1, OSRC and RCC4), normal kidney cell lines (HK-2 and 293) and primary cancer cells isolated from human ccRCC tissues. LDs are indicated by white arrows. Scale bar, 20 μm. **(b)** Quantitation of LD amount in primary cancer cells and cell lines, normalized by the LD amount in primary cancer cells. **(c)** Representative Raman spectra of LDs in cell lines (786-O, 769-P, Caki-1, OSRC and HK-2) and primary cancer cells. Spectral intensity was normalized by the peak at 1,442cm^-1^. The black arrow indicates the band of cholesterol rings at 702 cm^-1^ in primary cancer cells. **(d)** Quantitation of LD amount and CE percentage in primary cancer cells before and after cryopreservation. Each triangle represents LD amount for a primary cell and each circle represents CE percentage for a LD. LD amount was normalized by the primary cancer cells group before cryopreservation. **(e)** Significantly differential expression of genes involved in the cholesterol metabolic pathway in primary cancer cells before and after cryopreservation for three patients. **(f)** Quantitation of LD amount in primary cancer cells after cultured *in vitro* for 1, 3, 5, 7, 9, 11 days. Each triangle represents LD amount for a primary cell. LD amount was normalized by the primary cancer cells group after cultured *in vitro* at the first day. **(g)** Quantitation of CE percentage in primary cancer cells after cultured *in vitro* for 1, 3, 5, 7, 9, 11 days. **(h)** Quantitation of CE percentage for LDs with detectable CEs in primary cancer cells after cultured *in vitro* for 1, 3, 5, 7, 9, 11 days. Each circle represents CE percentage for a LD in **g** and **h**. Error bar represents SEM. *p < 0.05, **p < 0.005, ***p < 0.0005.

Although cryopreservation is a routine procedure for cell culture, both LD amount and CE level were significantly reduced in primary cells after thawing from cryopreservation (Fig. 2d, S6a, and S6b). To further investigate the reason why cryopreservation reduced accumulation of LDs and CEs, whole-transcriptome RNA sequencing was performed. As shown in Fig. 2e, the cholesterol transport related gene STARD3 and the cholesterol efflux related gene NR1H2 were upregulated, whereas the lipid regulation related genes PIK3CA and PIK3CB, the low-density lipoprotein (LDL) uptake related gene LRP2, and the CE hydrolysis related genes NCEH1 and LIPA were downregulated, after thawing from cryopreservation. Such alterations in gene expression imply that cryopreservation may reduce LD amount and CE level by enhancing cholesterol efflux and inhibiting cholesterol uptake.

In addition, our data showed a gradual reduction in both LD amount (Fig. 2f and S6c) and CE level (Fig. 2g) over time with prolonged culture *in vitro*. Particularly, the LD mount gradually decreased to around 70% and the average CE percentage gradually reduced around half on Day 7. As shown in Fig. 2g, the CE percentage for different LDs varied, that is, some of LDs contained high levels of CEs while the others might not have detectable CEs. Due to such variations, regarding the analysis of CE level in this study, we quantified not only the average CE percentage, but also the fraction of LDs with detectable CEs and the CE percentage of the LDs with detectable CEs. As shown in Fig. S6d, the fraction of LDs with detectable CEs decreased gradually from 100% on Day 1 to 50% on Day 7. Within the LDs with detectable CEs, the CE percentage did not change significantly until Day 11 (Fig. 2h).

Thus, to avoid the influence of cryopreservation and culture time, we only used primary cancer cells at the first subculture and compare cells with or without treatment on the same day. Collectively, primary cancer cells retain the CE-rich phenotype of ccRCC and thus can be used as an appropriate model for the study of CE accumulation in ccRCC.

### CE accumulation in ccRCC is induced by VHL mutation and subsequent upregulation of HIFα

Given that VHL is a prevalently mutated tumor suppressor in ccRCC ^31^, we collected primary ccRCC cells from 17 patients and divided them into VHL mutation group (n = 11) and VHL non-mutation group (n = 6) (Table S2). Although the LD amount was not significantly different between the primary cancer cells from patients with and without VHL mutation (Fig. 3a), the cells in VHL mutation group showed much stronger Raman peaks for CEs compared to those in VHL non-mutation group (Fig. 3b). Quantitatively, the CE level of primary cancer cells with VHL mutation was 68.4% ±4.3%, which was significantly higher than that in cells without VHL mutation (21.7% ±9.3%) (Fig. 3c). Particularly, primary cells from two out of the six patients without VHL mutation contained no detectable CEs (Fig. 3c). The detailed quantifications of LD amount and CE percentage for each patient with or without VHL mutation are shown in the Table S2. These findings indicate a potential link between VHL mutation and CE accumulation in ccRCC.

**Fig. 3:**
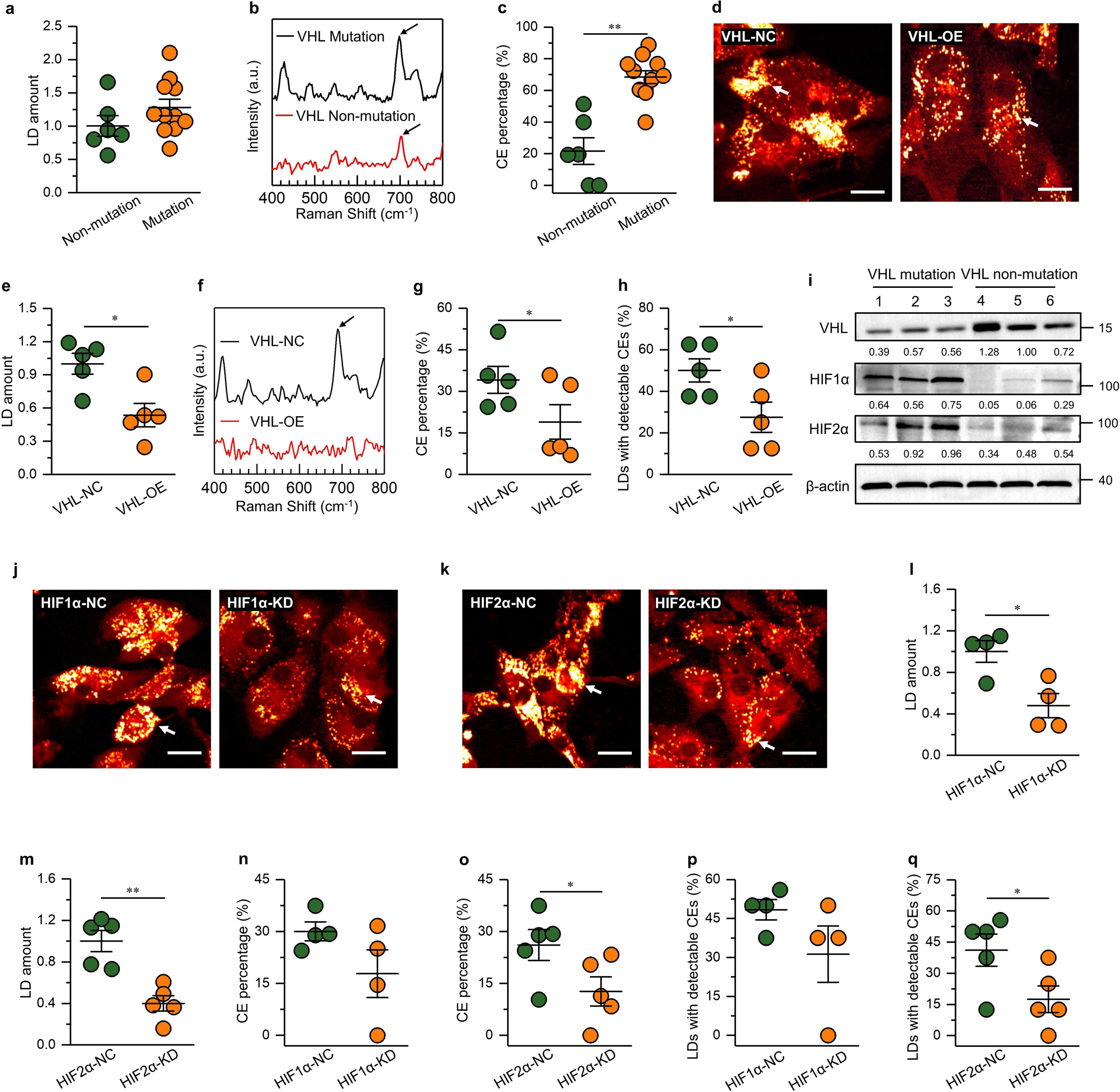
CE accumulation in ccRCC driven by VHL mutation and subsequent upregulation of HIFα. **(a)** Quantitation of LD amount in primary cancer cells with and without VHL mutation, normalized by the non-mutation group. **(b)** Representative Raman spectra of LDs in primary cancer cells with and without VHL mutation. Spectral intensity was normalized by the peak at 1,442 cm^-1^. The bands of cholesterol ring at 702 cm^-1^ are indicated by black arrows. **(c)** Quantitation of CE percentage in primary cancer cells with and without VHL mutation. **(d)** Representative SRS images of VHL-mutated primary cancer cells transfected with lentiviral negative control (VHL-NC) and VHL (VHL-OE). LDs are indicated by white arrows. Scale bar, 20 μm. **(e)** Quantitation of LD amount in primary cancer cells of the VHL-NC and VHL-OE groups. LD amount was normalized by the VHL-NC group. **(f)** Representative Raman spectra of LDs in primary cancer cells of the VHL-NC and VHL-OE groups. Spectral intensity was normalized by the peak at 1,442 cm^-1^. The band of cholesterol ring at 702 cm^-1^ is indicated by black arrow. **(g)** Quantitation of CE percentage in primary cancer cells of the VHL-NC and VHL-OE groups. **(h)** The fraction of LDs with detectable CEs in primary cancer cells of the VHL-NC and VHL-OE groups. **(i)** Immunoblot of antibodies against VHL, HIF1α, HIF2α, and β-actin in 6 independent cases of human ccRCC tissues with VHL mutation (n = 3) or VHL non-mutation (n = 3). **(j)** Representative SRS images of VHL-mutated primary cancer cells stably transfected with negative control (HIF1α-NC) and HIF1α shRNA (HIF1α-KD). LDs are indicated by white arrows. Scale bar, 20 μm. **(k)** Representative SRS images of VHL-mutated primary cancer cells stably transfected with negative control (HIF2α-NC) and HIF2α shRNA (HIF2α-KD). LDs are indicated by white arrows. Scale bar, 20 μm. **(l, m)** Quantitation of LD amount in primary cancer cells of the HIF1α-NC/HIF2α-NC and HIF1α-KD/HIF2α-KD groups. LD amount was normalized by the HIF1α-NC or HIF2α-NC group. **(n, o)** Quantitation of CE percentage in primary cancer cells of the HIF1α-NC/HIF2α-NC and HIF1α-KD/HIF2α-KD groups. **(p, q)** The fraction of LDs with detectable CEs in primary cancer cells of the HIF1α-NC/HIF2α-NC and HIF1α-KD/HIF2α-KD groups. Each circle represents average LD amount for one patient in **a**, **e**, **l**, and **m**. Each circle represents average CE percentage for one patient in **c**, **g**, **n** and **o**. Each circle represents the average fraction of LDs with detectable CEs for one patient in **h**, **p** and **q**. Error bars represent SEM. *p < 0.05, **p < 0.005.

To further determine whether VHL mutation was involved in CE accumulation, wild-type VHL was stably reintroduced into the primary cancer cells with VHL mutation, which resulted in significant reductions in both LD amount (Fig. 3d and 3e) and CE level (Fig. 3f-3h). Specifically, there was a remarkable reduction in the Raman peaks for CEs upon VHL overexpression (Fig. 3f). Quantitatively, the average CE percentage was significantly reduced in VHL overexpression group compared to the control group (Fig. 3g). Furthermore, the fraction of LDs with detectable CEs was found to be significantly decreased upon VHL overexpression (Fig. 3h), while the CE level of the LDs with detectable CEs remained unchanged (Fig. S7a). Detailed information regarding LD amount and CE level for each patient is shown in Fig. S7b, S7c, Table S3, and S4. Together, these results demonstrate that CE accumulation is likely driven by VHL mutation in ccRCC.

VHL mutation leads to degradation failure and constitutive stabilization of HIFα, which is considered as the major driving force for ccRCC development ^11,12^. We examined the protein expression levels of VHL, HIF1α, and HIF2α in the clinical samples. As shown in Fig. 3i, the results revealed that HIF1α and HIF2α expression levels were markedly upregulated in VHL mutated ccRCC tissues compared to those without VHL mutation. Further western blot data confirmed the reduction in HIFα expression upon VHL overexpression in primary cancer cells (Fig. S7d). Although HIFα has been shown to regulate fatty acid metabolism ^8,9^, its role in CE accumulation remains elusive. To address this question, we tested if knockdown of HIFα by shRNA could reduce CE accumulation in primary cancer cells. As shown in Fig. 3j and 3k, LD accumulation in primary cancer cells was obviously decreased upon knockdown of either HIF1α or HIF2α. Quantitatively, the LD amount was reduced about half in the cells of both HIF1α knockdown group and HIF2α knockdown group (Fig. 3l and 3m). In terms of the CE level in primary cancer cells, the average CE percentage was not significantly changed upon HIF1α knockdown (Fig. 3n), but significantly reduced upon HIF2α knockdown (Fig. 3o). Consistently, the fraction of LDs with detectable CEs was not changed upon HIF1α knockdown (Fig. 3p), but significantly reduced upon HIF2α knockdown (Fig. 3q). The CE level of the LDs with detectable CEs remained unchanged in both HIF1α knockdown group and HIF2α knockdown group (Fig. S7e and S7f). Detailed information regarding LD amount and CE level for each patient is shown in Fig. S7g, S7h, Table S3, S5, and S6. Together, these results suggest that CE accumulation is probably induced by upregulation of HIFα, mainly HIF2α.

### CE accumulation in ccRCC is dependent on the upregulation of PI3K/AKT/mTOR/ SREBP pathway

VHL and HIFα have been found to be closely related with PI3K/AKT/mTOR signaling pathway in ccRCC^32^. Our western blot data confirmed that both VHL overexpression and HIFα knockdown induced significant reductions in the expression of p-AKT and p-S6 (Fig. S7d, S7i, and S7j). As one of the interfaces of oncogenic signaling and cancer metabolism, PI3K/AKT/mTOR pathway is known to promote ccRCC ^33^, but its link with CE accumulation is not clear. To clarify this question, we inhibited the PI3K/AKT/mTOR pathway in primary cancer cells with LY294002 (a selective PI3K inhibitor), MK2206 (a selective AKT inhibitor), and rapamycin (a selective mTOR inhibitor), respectively (Fig. 4a). As shown in Fig. 4b and 4c, inhibition of PI3K/AKT/mTOR pathway significantly reduced LD amount in primary cancer cells. Regarding the CE level, although the average CE percentage was not significantly reduced due to big variations among patients (Fig. 4d), the fraction of LDs with detectable CEs was significantly decreased by PI3K/AKT inhibition in primary cancer cells (Fig. 4e). Detailed information regarding LD amount and CE level for each patient is shown in Fig. S8a, S8b, Table S3, and S7.

**Fig. 4:**
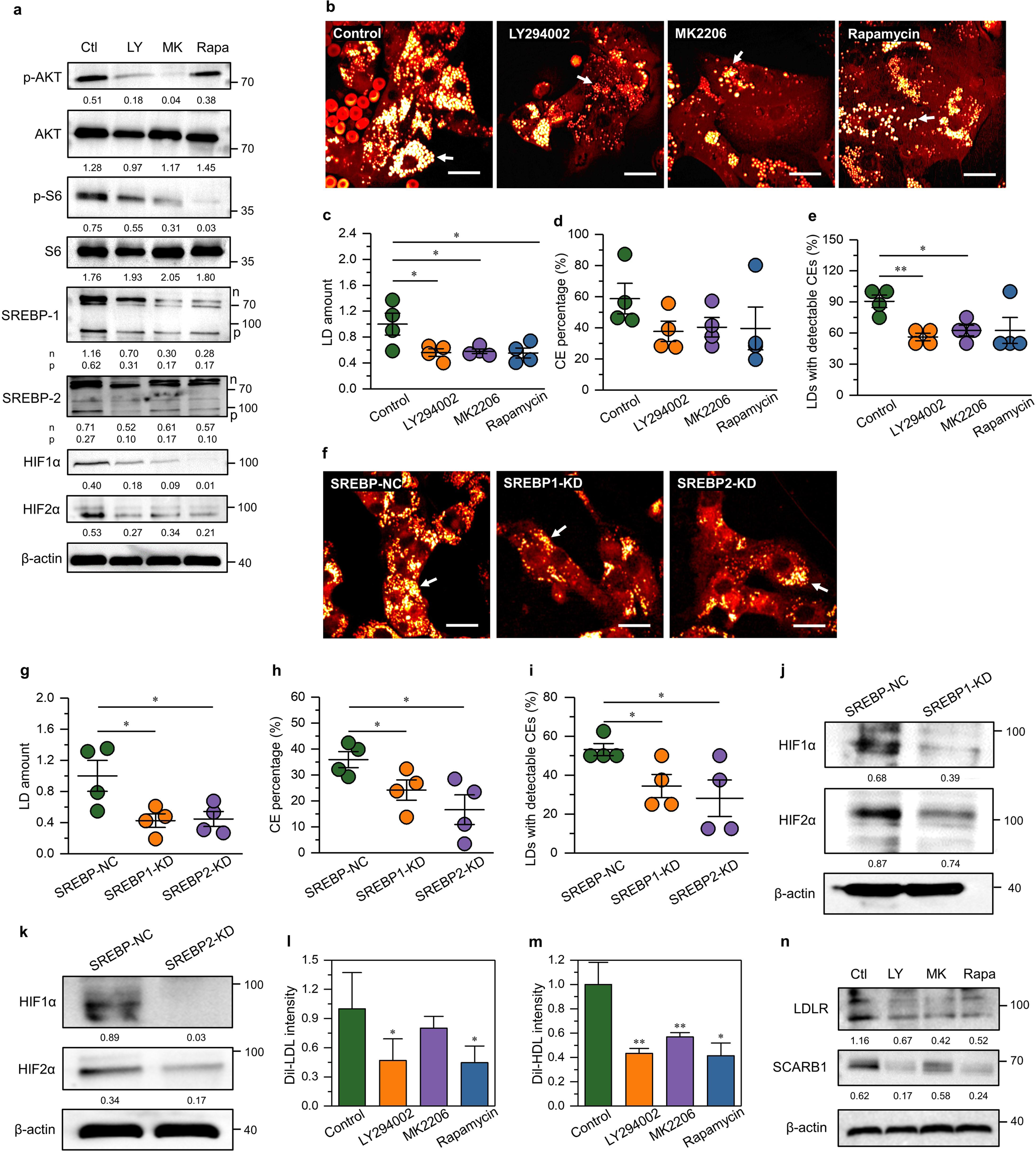
CE accumulation in ccRCC induced by the upregulation of PI3K/AKT/mTOR/SREBP pathway. **(a)** Immunoblot of antibodies against p-AKT, AKT, p-S6, S6, SREBP-1, SREBP-2, HIF1α, HIF2α, and β-actin in primary cancer cells treated with DMSO as control (Ctl), LY294002 (50 μM, 3 day) (LY), MK2206 (10 μM, 2 day) (MK), and rapamycin (100 nM, 2 day) (Rapa). p-SREBP, SREBP precursor; n-SREBP, nuclear SREBP. **(b)** Representative SRS images of primary cancer cells of control, LY294002, MK2206, and rapamycin groups. LDs are indicated by white arrows. Scale bar, 20 μm. **(c)** Quantitation of LD amount in primary cancer cells of control, LY294002, MK2206, and rapamycin groups. LD amount was normalized by the control group. **(d)** Quantitation of CE percentage in primary cancer cells of control, LY294002, MK2206, and rapamycin groups. **(e)** The fraction of LDs with detectable CEs in primary cancer cells of control, LY294002, MK2206, and rapamycin groups. **(f)** Representative SRS images of primary cancer cells transfected with negative control (SREBP-NC), SREBP-1 shRNA (SREBP1-KD), and SREBP-2 shRNA (SREBP2-KD). LDs are indicated by white arrows. Scale bar, 20 μm. **(g)** Quantitation of LD amount in primary cancer cells of the SREBP-NC, SREBP1-KD, or SREBP2-KD groups. LD amount was normalized by the SREBP-NC group. **(h)** Quantitation of CE percentage in primary cancer cells of the SREBP-NC, SREBP1-KD, and SREBP2-KD groups. **(i)** The fraction of LDs with detectable CEs in primary cancer cells of the SREBP-NC, SREBP1-KD, and SREBP2-KD groups. Each circle represents average LD amount for one patient in **c** and **g**. Each circle represents average CE percentage for one patient in **d** and **h**. Each circle represents the average fraction of LDs with detectable CEs for one patient in **e** and **i**. Error bars represent SEM. *p < 0.05, **p < 0.005. **(j, k)** Immunoblot of antibodies against HIF1α, HIF2α, and β-actin in primary cancer cells of the SREBP-NC, SREBP1-KD, and SREBP2-KD groups. **(l, m)** Quantitation of DiI-LDL or Dil-HDL uptake in primary cancer cells treated with DMSO as control, LY294002 (50 μM, 3 day), MK2206 (10 μM, 2 day), and rapamycin (100 nM, 2 day) (n = 3 patients). DiI-LDL/HDL intensity was normalized by the control group. **(n)** Immunoblot of antibodies against LDLR, SCARB1, and β-actin in primary cancer cells treated without (Ctl) or with LY294002 (LY), MK2206 (MK), and rapamycin (Rapa).

Sterol regulatory element-binding proteins (SREBPs), which can be controlled by PI3K/AKT/mTOR, mainly regulate genes involved in intracellular lipid homeostasis ^34^. Our western blot data confirmed that the inhibition of PI3K/AKT/mTOR induced downregulation of both SREBP-1 and SREBP-2 (Fig. 4a, Fig. S8c, and Fig. S8d). To figure out which specific isoform of SREBPs was affected, the mRNA levels of three SREBP isoforms, including SREBP-1a, SREBP-1c, and SREBP-2 were measured via qPCR. As shown in Fig. S8e, the inhibition of the PI3K pathway significantly reduced expressions of SREBP-1a and SREBP-2. Subcequently, the target genes of SREBPs for cholesterol synthesis and uptake (HMGCS1, LDLR) and fatty acid synthesis (FASN, SCD1) were also significantly inhibited. Upon knockdown of SREBP-1 and SREBP-2 (Fig. S8f and S8g), the LD amount, the average CE percentage, and the fraction of LDs with detectable CEs were all significantly decreased in primary cancer cells (Fig. 4f-4i). Notably, the knockdown of SREBP-2, which is specifically responsible for regulating cholesterol synthesis and uptake, affected the CE level more evidently (Fig. 4h and 4i). Detailed information regarding LD amount and CE level for each patient is shown in Fig. S5c, S5d, Table S3 and S8. The reduced expressions of the SREBP target genes upon knockdown of SREBP-1 and SREBP-2 are shown in Fig. S8h and S8i. Moreover, inhibition of the PI3K/AKT/mTOR pathway effectively decreased the uptake of DiI-labeled LDL (Fig. 4l) and DiI-labeled HDL (Fig. 4m), which were consistent with the reduced expression levels of LDLR and SCARB1, respectively (Fig. 4n). In addition, the western blots revealed that inhibition of the PI3K/AKT/mTOR pathway and SREBPs in turn suppressed the HIFα expression (Fig. 4a, 4j, and 4k). Together, these results reveal that CE accumulation in ccRCC is promoted by the PI3K/AKT/mTOR/SREBP pathway, which shows a positive feedback loop with HIFα.

### CE accumulation relied on cholesterol esterification in ccRCC

Excess free cholesterol needs to be converted to CEs by sterol O-acyltransferase (SOAT) enzymes and then stored into LDs to avoid toxicity ^35^. As excepted, inhibition of SOAT enzymes with a potent SOAT inhibitor, avasimibe, induced significant reductions in LD amount, the average CE percentage, and the fraction of LDs with detectable CEs in primary cancer cells (Fig. 5a-5c). Detailed information regarding LD amount and CE level for each patient is shown in (Fig. S9a, S9b, Table S3, and S9). Collectively, these results demonstrate that CE accumulation in ccRCC relies on sufficient cholesterol esterification.

**Fig. 5:**
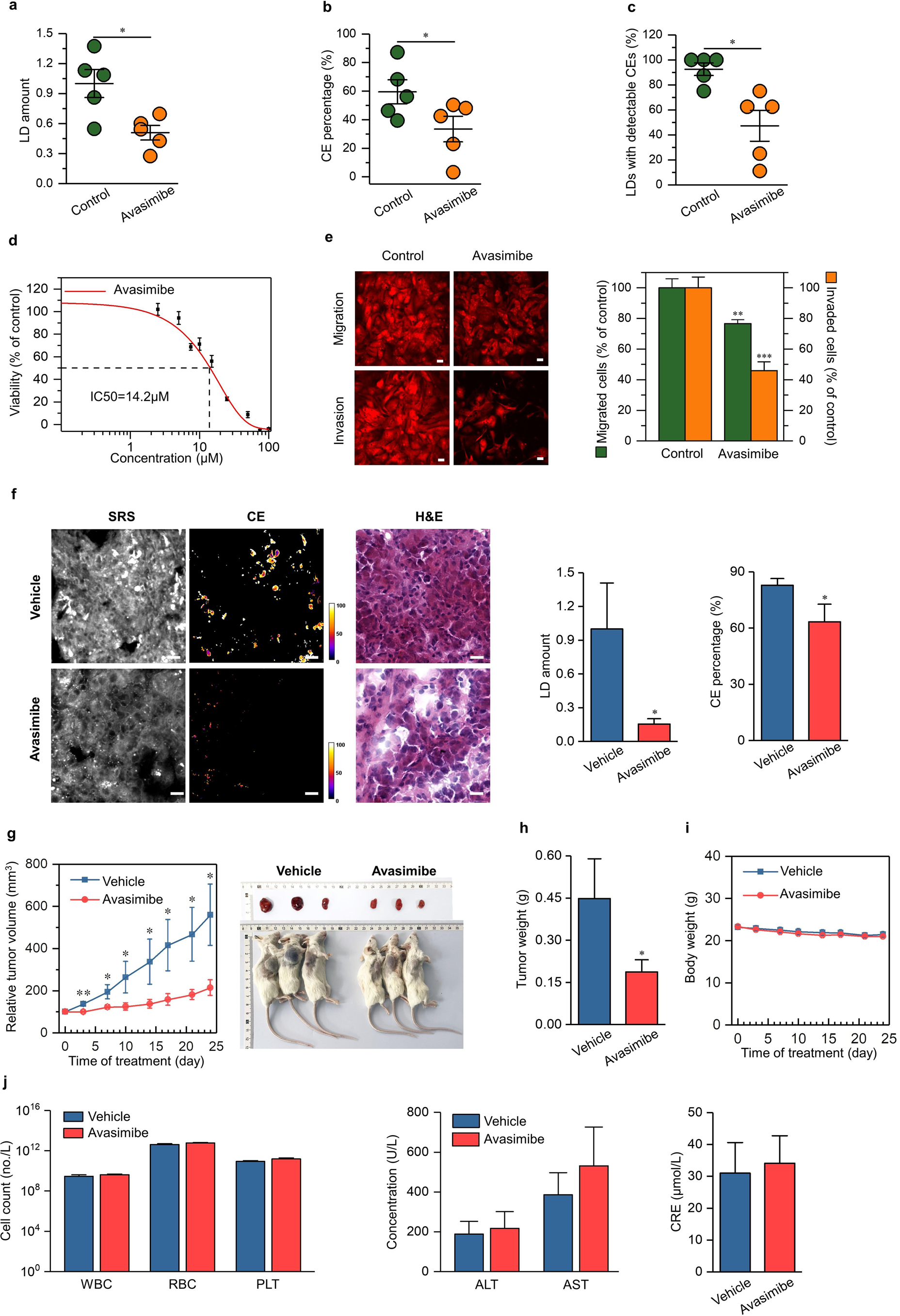
Inhibition of cholesterol esterification impairs ccRCC aggressiveness *in vitro* and *in vivo*. **(a)** Quantitation of LD amount in ccRCC primary cells treated with DMSO as control and avasimibe (20 μM, 2 day). LD amount was normalized by the control group. Each circle represents average LD amount for one patient. **(b)** Quantitation of CE percentage in primary cancer cells of control and avasimibe groups. Each circle represents average CE percentage for one patient. **(c)** The fraction of LDs with detectable CEs in primary cancer cells of control and avasimibe groups. Each circle represents the average fraction of LDs with detectable CEs for one patient. **(d)** IC50 curve of primary cancer cells upon avasimibe treatment (3 day) (IC50 ≈ 15 μM). **(e)** Representative images and quantitation of migration and invasion of primary cancer cells upon avasimibe treatment (15 μM, 3 day) (n = 7). The control group (DMSO) was used for normalization. **(f)** Representative images of hyperspectral SRS (at 2850 cm^-1^), CE percentage and H&E staining (left), quantitation of LD amount (middle), and CE percentage (right) of tumor tissues harvested from the vehicle and avasimibe-treated PDX mice (n = 6). LD amount was normalized by the control group. Scale bar, 20 μm. **(g)** Relative tumor volume (left) and representative images (right) of PDX mice treated daily with vehicle and avasimibe (15mg/kg) (n = 10). **(h)** The weight of tumor tissues harvested from the vehicle and avasimibe-treated PDX mice on day 24 (n = 10). **(i)** Body weight of the vehicle and avasimibe-treated PDX mice (n = 10). **(j)** Hematological and blood biochemical analysis in the vehicle and avasimibe-treated PDX mice on day 24 (n = 10). WBC, white blood cell; RBC, red blood cell; PLT, platelet; ALT, alanine aminotransferase; AST, aspartate aminotransferase; CRE, creatinine. Error bars represent SEM. *p < 0.05, **p < 0.005, ***p < 0.0005.

### CE depletion impaired ccRCC aggressiveness *in vitro* and *in vivo*

Considering that aberrant CE accumulation occurs in ccRCC but not normal kidney, here we explored whether cholesterol esterification affected ccRCC viability, migration, and invasion. Firstly, CE depletion by avasimibe significantly reduced viability of primary cancer cells with IC_50_ value of ∼15 μM (Fig. 5d, S9e, and S9f), while the inhibitory effects by avasimibe were much more moderate in the commonly used ccRCC cell lines (786-O, 769-P, Caki-1, OSRC, and RCC4) and normal kidney cell line (HK-2) (Fig. S9g). Further flow cytometry analysis revealed that avasimibe treatment significantly prohibited the transition from G1 to S phase of primary cancer cells (Fig. S9h), which might explain why avasimibe reduced cell viability. Secondly, results of the Transwell experiments indicated that CE depletion by avasimibe significantly suppressed migration and invasion capabilities of primary cancer cells (Fig. 5e). These results suggest that CE depletion substantially suppresses ccRCC aggressiveness *in vitro*.

To investigate the anti-cancer effect of avasimibe *in vivo*, we further developed a subcutaneous patient-derived tumor xenograft (PDX) mouse model of ccRCC. The spectral imaging data confirmed that the PDX model could retain the CE-rich phenotype of human ccRCC (Fig. 5f). As shown in Fig. 5 and S10, daily avasimibe treatment for 24 days remarkably suppressed tumor volume by ∼62% (Fig. 5g and S10a) and tumor weight by ∼58% (Fig. 5h). Spectroscopic analysis of extracted tissues revealed significant reductions in both LD amount (by ∼84%) and CE level (by ∼18%) (Fig. 5f) of avasimibe-treated mice compared with the vehicle group, indicating that avasimibe worked to inhibit CE formation in tumor cells *in vivo*. Immunofluorescence using markers for apoptosis (TUNEL) showed that avasimibe significantly increased apoptosis (Fig.S10b).

Importantly, the avasimibe treatment did not cause general toxicity to the mice. First, no significant changes in body weight were observed in the mice treated with avasimibe (Fig. 5i). Second, pathological review of sections of liver, lung, heart, spleen, kidney, and adrenal gland harvested from mice receiving avasimibe showed no detectable signs of toxicity (Fig. S10c). Third, the hematological results, including the number of white blood cell (WBC), red blood cell (RBC), and platelet (PLT), had no significant changes in avasimibe treated mice (Fig. 5j). Fourth, the biochemical analysis of the levels of alanine aminotransferase (ALT), aspartate aminotransferase (AST), and creatinine (CRE) revealed that avasimibe did not damage the liver function or kidney function.

Subsequently, we conducted a pharmacokinetic study of avasimibe treatment by using liquid chromatography-mass spectroscopy (LC-MS). The plasma concentration profile of avasimibe after intraperitoneal administration was shown in Fig. S10d, and the detailed pharmacokinetic parameters were presented in Table S10. Avasimibe quickly reached its maximum plasma concentration at 0.5 hour with a Cmax of 18.8 μg/mL, and decreased gradually over 24 hours. Next, to evaluate distribution of avasimibe in tissues, tumor, liver, lung, heart, spleen, kidney, brain, adrenal gland, intestine, muscle, bone, and feces samples were collected at 0.5 hour (Tmax) post administration. As shown in Fig. S10e, avasimibe was detected widely in most tissues, including tumors. Collectively, these results show that avasimibe is a promising candidate against ccRCC owing to its superior therapeutic effect, negligible toxicity, and good pharmacokinetic characteristics.

### CE depletion impaired ccRCC aggressiveness through downregulation of the integrin and MAPK signaling pathways

To elucidate the underlying mechanisms by which CE depletion impaired ccRCC aggressiveness, we firstly analyzed differentially expressed genes in primary cancer cells treated with avasimibe using whole-transcriptome RNA sequencing (Fig. 6a). In consistency with the inhibitory effects of avasimibe, various genes related to cellular processes were significantly downregulated in the avasimibe-treated group compared with the control group. Importantly, multiple genes related to the integrin pathway (e.g., ITGA6 and ITGB1), which plays a crucial role in cancer progression, were significantly downregulated upon avasimibe treatment. The western blot data and quantitative confocal microscopic analyses also confirmed the reduced expressions of ITGA6 and ITGB1 in the avasimibe-treated group (Fig. 6b and 6c). Given that integrins are located on the plasma membrane and regulated by the membrane cholesterol level ^36^, we further clarified how membrane cholesterol was affected by avasimibe in primary cancer cells. As shown in Fig. 6d and 6e, the fluorescence intensity of filipin, which is commonly used to map intracellular free cholesterol, was remarkedly decreased on cell membranes of primary cancer cells treated with avasimibe. Further biochemical assay confirmed significantly reduced free cholesterol level on cell membranes of the avasimibe-treated group (Fig. 6f).

**Fig.6:**
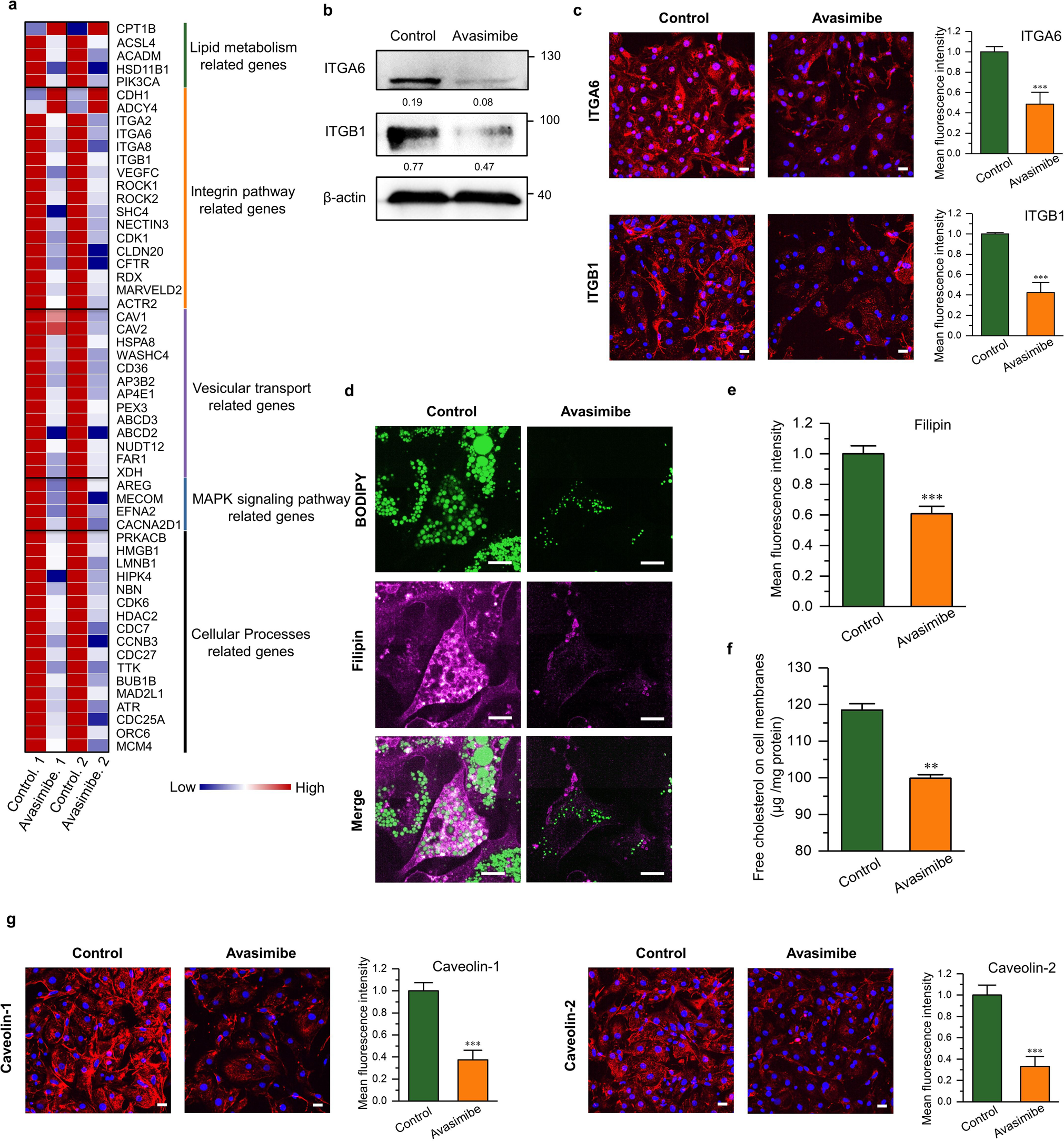
CE Depletion Reduces ccRCC aggressiveness via downregulation of the integrin and MAPK signaling pathways. **(a)** Differential gene expression of primary cancer cells upon avasimibe treatment (20 μM, 2 day) for two independent patients. **(b)** Immunoblot of antibodies against ITGA6, ITGB1, and β-actin in primary cancer cells of control and avasimibe groups. **(c)** Representative images and quantification of mean fluorescence intensity of ITGA6 and ITGB1 staining in primary cancer cells upon avasimibe treatment. Scale bar, 20 μm. **(d)** Representative images of filipin staining for free cholesterol ( magenta color) and BODIPY staining for LDs (green color) in primary cancer cells upon avasimibe treatment. Scale bar, 5 μm. **(e)** Quantification of mean fluorescence intensity of filipin staining in **d**. **(f)** Free cholesterol levels on cell membranes in primary cancer cells upon avasimibe treatment (n = 3). **(g)** Representative images and quantification of mean fluorescence intensity of Caveolin-1 and Caveolin-2 staining in primary cancer cells upon avasimibe treatment. The mean fluorescence intensity was normalized to the control group, respectively (n ≥ 6) in **c**, **e**, and **g**. Scale bar, 20 μm. Error bars represent SEM. *p < 0.05, **p < 0.005, ***p < 0.0005.

Besides the integrin pathway, caveolin-1 (CAV1) and several genes associated with MAP kinase (MAPK) signaling pathway were also markedly downregulated in primary cancer cells upon avasimibe treatment (Fig. 6a). The quantitative confocal microscopic analyses also confirmed the reduced expressions of caveolins in the avasimibe-treated group (Fig. 6g). As a regulator of intracellular cholesterol homeostasis, CAV1 has been shown to be widely upregulated in various cancers and promote tumor progression and drug resistance ^37^. CAV1-mediated MAPK pathway may contribute to cancer cell survival and metastasis ^22,38^. Taken together, these results suggest that CE depletion impaired ccRCC aggressiveness due to reduced cholesterol levels on cell membranes and subsequent downregulation of integrin and MAPK signaling pathways.

## Discussion

Although CE accumulation has been recognized as a hallmark of ccRCC, its exact role in ccRCC progression remains elusive. As illustrated in Fig. 6, our study unraveled that CE accumulation in LDs was induced by the mutation of tumor suppressor VHL and subsequent activation of the HIFα mediated PI3K/AKT/mTOR pathway. Inspiringly, inhibition of cholesterol esterification effectively suppressed ccRCC progression via reduced membrane cholesterol levels and the subsequent downregulation of integrin and MAPK signaling pathways. As discussed below, these findings improve current understanding of the molecular mechanisms and pathological significance of CE accumulation in ccRCC progression, which may provide a potential metabolic target for ccRCC treatment.

Firstly, our study demonstrates the need for primary cancer cell, instead of cell lines, as the suitable model to study CE accumulation in ccRCC. Enabled by the label-free Raman spectromicroscopy platform, which integrated stimulated Raman scattering microscopy with confocal Raman spectroscopy, we could quantitatively analyze LD distribution and composition at the single cell level in intact cancer cell and tissue specimens *in situ* without any processing or exogenous labeling. Our results revealed that, in contrast with the high CE level in human ccRCC tissues, the commonly used ccRCC cell lines in literatures did not contain detectable CEs in LDs. Thus, it is not surprising that most previous mechanistic studies about LD accumulation in ccRCC focused on the altered metabolism of fatty acid but not cholesterol ^8–10^. This is very likely the culprit for current limited understanding of CE accumulation in ccRCC. In order to address this problem, we isolated primary cancer cells from human ccRCC tissues and showed that they could represent the signature of ccRCC with CE level as high as the original tissue. Collectively, our study indicates that primary cancer cell is an appropriate cell model to elucidate the role of CE in ccRCC.

Secondly, our study extends the current understanding of cholesterol metabolism in ccRCC. As the most common mutation of ccRCC, tumor suppressor VHL mutation induces constitutive stabilization of HIFα ^11,12^, which has been shown to promote the storage of fatty acids in the form of triacylglycerol in LDs ^8^. Nevertheless, the role of VHL and HIFα in the regulation of cholesterol metabolism in ccRCC has not been investigated. Our results unraveled that aberrant CE accumulation in ccRCC was a consequence of VHL mutation and subsequent activation of HIFα, which has not been documented before. DNA methylation alteration is another cause of VHL loss in renal cancer^39^. In this work, we focus on the altered cholesterol metabolism in ccRCC with VHL mutation, which does not include DNA methylation. The effect of VHL methylation on lipid metabolism in ccRCC is another important topic, which deserves further investigation in future work. Moreover, our study elucidates how VHL/HIFα regulates CE accumulation in ccRCC. Upregulations of PI3K/AKT/mTOR signaling pathway and subsequent SREBPs have been shown to alter lipid metabolism in a variety of human cancers ^1^. Thus, it is intriguing to explore whether VHL/HIFα links to CE accumulation via the PI3K/AKT/mTOR/SREBP pathway. Our results uncovered that VHL mutation and subsequent stabilization of HIFα, mainly HIF2α, upregulated the PI3K/AKT/mTOR/SREBP pathway, which induced CE accumulation. Actually, HIF1α and HIF2α share several common transcriptional targets, but in the meanwhile exhibit unique functions at different stages of tumor formation and progression ^40,41^. In common, both HIF1α and HIF2α have been proved to have positive correlations with the PI3K/AKT/mTOR/SREBPs pathway ^42^. Notably, HIF1α is mainly responsible for glycolysis whereas HIF2α regulates lipid metabolism, ribosome biogenesis, angiogenesis, and so on ^40,43^. As reported before, the activation of HIF2α, rather than HIF1α, in VHL-deficient tumor cells increases mTORC1 activity and consequently promotes synthesis of cholesterol through SREBP-1a and SREBP-2 ^44,45,46^. This partly explains why HIF2α was found to be the major player in CE accumulation in ccRCC. Furthermore, VHL/HIF and PI3K/AKT/mTOR/SREBPs pathways interact in a complex signaling network. VHL mutation and subsequent stabilization of HIFα upregulate the PI3K/AKT/mTOR pathway through various growth factors and their corresponding transmembrane receptor tyrosine kinases ^42,47^. The subsequent regulated SREBPs directly activate the cholesterol biosynthesis ^48^, which is a highly oxygen-intensive process ^49^. This may be an important reason for HIFα upregulation upon SREBP activation. Thus, VHL/HIF and PI3K/AKT/mTOR/SREBPs pathways form a complex signaling network with a positive feedback loop contributing to intracellular CE accumulation and ccRCC development.

Regarding the source of cholesterol for CE accumulation in ccRCC, we found that exogenous lipoprotein uptake was the primary source of cholesterol for CE accumulation in ccRCC. Our data showed that both LDL uptake via LDLR and HDL uptake via SCARB1 contributed to CE accumulation in ccRCC and regulated by the PI3K/Akt/mTOR pathway (Fig. S11a-S11d and Fig. 4l-4n). Additionally, expression of SCARB1, but not LDLR, was greatly elevated in human ccRCC tissues compared with normal kidney tissues (Fig. S11e), and also higher in ccRCC primary cell relative to cell lines (Fig. S11f and S11g). The findings about the greater dependence on HDL uptake as the source of cholesterol in ccRCC are consistent with the previous literature^15^.

Finally, our study discovers cholesterol esterification as a vulnerable metabolic target for ccRCC treatment. Although surgery is considered as the first choice for ccRCC treatment, adjuvant therapies are still crucial to reduce recurrence and prolong survival. Unfortunately, ccRCC lacks sensitivity to chemotherapy and often develops resistance to current targeted therapies against vascular endothelial growth factor (VEGF) and mTOR^6^. Recently developed immunotherapy could extend survival, but only benefit a limited population of patients ^6^. Thus, there is an urgent need to discover new therapeutic target for ccRCC. Our results showed that inhibition of cholesterol esterification by avasimibe, the potent inhibitor of SOAT, significantly impaired growth, migration and invasion of primary ccRCC cells, but not normal cells. Moreover, it is worth to mention that avasimibe was previously used to treat atherosclerosis and showed minimal toxicity ^50^. Avasimibe has also been used to effectively suppressed development and progression of many other types of cancers, such as prostate cancer, pancreatic cancer, glioblastoma, and liver cancer ^22,23,51,52^, which suggests that cholesterol esterification could become a universal therapeutic target for human cancers. Notably, the mechanism by which CE depletion impairs cancer progression varies in different cancers. Our results unraveled that inhibition of cholesterol esterification disturbed intracellular cholesterol homeostasis and reduced membrane cholesterol levels. According to the literatures^53–55^ that define three pools of membrane cholesterol, including PFO-accessible pool that is accessible to PFO binding, sphingomyelin (SM)-sequestered pool that binds PFO only after SM is destroyed by sphingomyelinase, and residual pool that does not bind PFO even after sphingomyelinase treatment, when lipoprotein-derived cholesterol is liberated in lysosomes, the PFO-accessible pool on the cell membrane is expanded, and after a short lag, the ER’s PFO-accessible regulatory pool is then increased. This regulatory mechanism allows cells to ensure optimal cholesterol levels in cell membrane while avoiding cholesterol overaccumulation. In this study, it was found that cholesterol esterification inhibition by avasimibe significantly reduced the uptake of LDL and HDL (Fig. S11h and S11i), and also expressions of the corresponding receptors LDLR and SCARB1 (Fig. S11j). Accordingly, less lipoprotein-derived cholesterol liberated may lead to reduced PFO-accessible pool on the cell membrane, which may partly explain the reduced free cholesterol level on cell membranes upon avasimibe treatment. Such membrane cholesterol reduction further led to downregulations of integrin signaling pathway and caveolin mediated MAPK pathway, both of which have been shown to promote cancer progression ^22,38^. Taken together, our study sheds new light on the mechanism underlying CE accumulation in ccRCC and opens up a new avenue for ccRCC treatment.

## Contributors

S.Y. and L.Y. designed and supervised the study. S.Z. and S.Y. developed the methodology. S.Z, Y.X.H, Y.Y.Z, Z.J.L., J.L., J.Y., A.B.H., S.H., Y.Q.G., Z.S.H., K.W.Y., Z.J.X., and W.Y. performed the experiments. T.F., S.Z, W.C.F., S.Y., and L.Q.Z. interpreted the data. S.Z, S.Y., and L.Y. wrote the manuscript. All authors critically reviewed the manuscript.

## Data sharing statement

All data associated with this study are available within the paper or the Supplementary Materials. Further information and requests for resources and reagents should be directed to the corresponding author.

## Supporting information

Zhang et al_CE in ccRCC_Supplementary

---

**Fig. 7:**
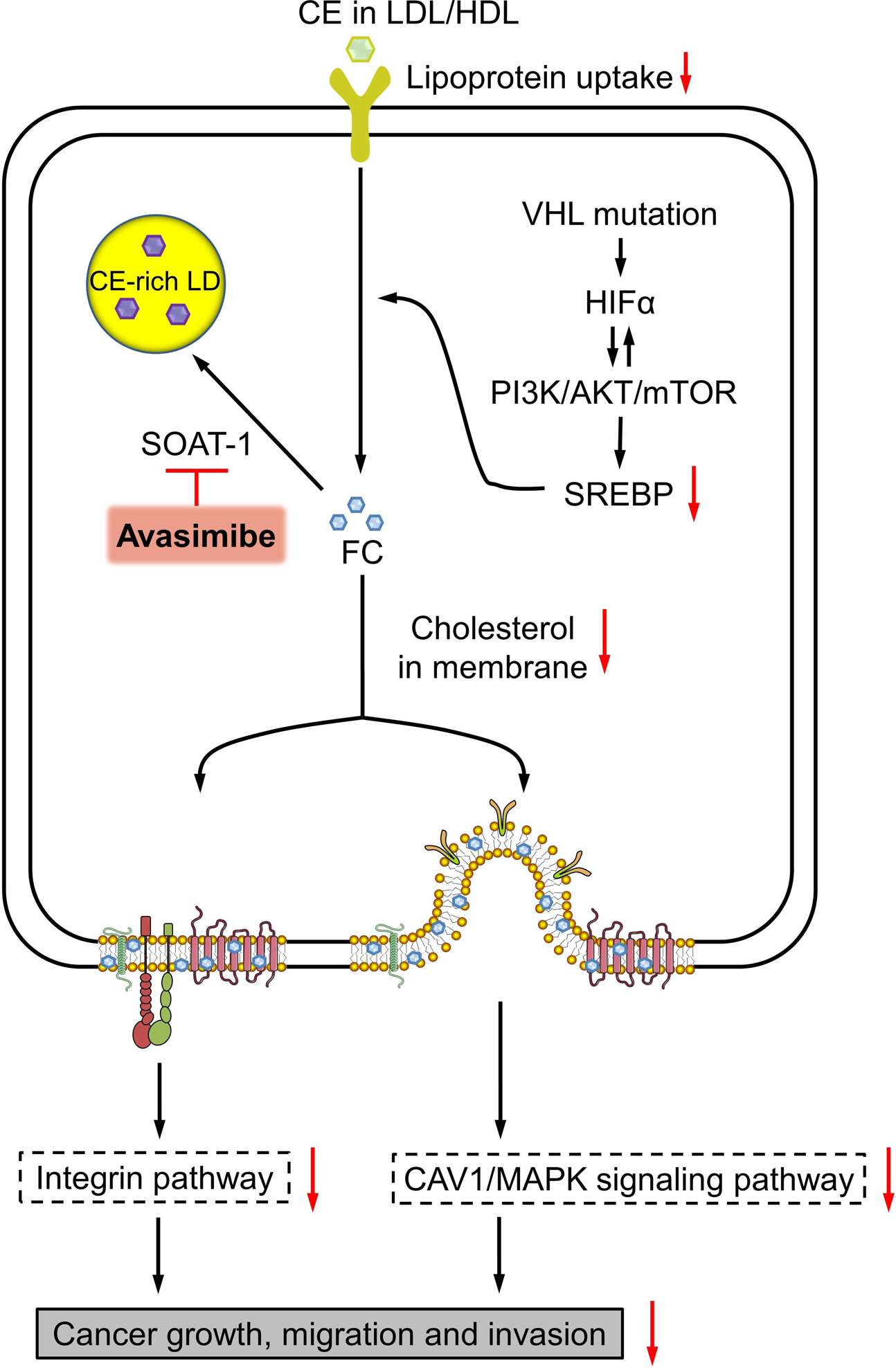
Molecular mechanism underlying CE accumulation in ccRCC and suppression of cancer progression upon CE depletion. The schematic shows that VHL mutation activates PI3K/AKT/mTOR/ SREBPs pathway in a HIFα-dependent manner. The excess free cholesterol (FC) is then esterified into CEs by SOAT-1 and stored into LDs. The red arrows indicate the consequences of CE depletion by avasimibe treatment. Inhibition of cholesterol esterification disturbs the cholesterol homeostasis by downregulations of SREBPs and lipoprotein uptake. Consequently, the reduced cholesterol level on cell membrane leads to downregulations of integrin and MAPK signaling pathways that further impair ccRCC progression.

