## Supplementary material for "VHL Mutation Drives Human Clear Cell Renal Cell Carcinoma Progression Through PI3K/AKT-Dependent Cholesteryl Ester Accumulation": Zhang et al_CE in ccRCC_Supplementary

### Supplementary Experimental Procedures

#### Human tissue specimens

All tissues specimens were obtained from ccRCC patients undergone nephrectomy at Peking University First Hospital. This study was approved by the Institutional Review Boards of both Beihang University (BM20220161) and Peking University First Hospital (2020keyan400). Informed written consent from all participants was obtained prior to the research. ccRCC tissues (n=24) and normal adjacent tissues (n=24) were obtained from ccRCC patients (n=41) undergone nephrectomy at Peking University First Hospital. Tissue specimens were snap-frozen in liquid nitrogen within 10 minutes of surgery excision. Each sample was ~100 mg. Frozen tissue samples were embedded in Optimal Cutting Temperature (OCT) media for further sectioning into pairs of neighboring slices, with one unstained 20-µm slice for spectroscopic imaging and the other neighboring 7-µm slice for H&E staining. Pathological examination was performed by experienced pathologists. The rest tissue samples were stored in -80°C refrigerator. Additionally, another 34 fresh ccRCC tissues were collected from independent patient cohort and put into cold DMEM/F12 medium on ice right after surgery excision for further isolation of primary cancer cells.

#### Cell cultures

##### *Cell lines*

Human ccRCC cell lines (786-O, 769-P, and Caki-1), normal proximal tubular cell line (HK-2) and normal embryonic kidney cell line (293) were obtained from the American Type Culture Collection. OSRC and RCC4 were provided by Dr. Anbang He (Peking University First Hospital). 786-O, 769-P and OSRC were cultured in RPMI 1640 (Gibco, cat 11875093) supplemented with 10% fetal bovine serum (FBS) (Biological Industries, cat 04-001-1ACS) and 1% penicillin/streptomycin (PS) (Gibco, cat 15070063). Caki-1 was cultured in McCoy's 5A (modified) medium (Gibco, cat 16600082) supplemented with 10% FBS and 1% PS. RCC4 and 293 were cultured in DMEM (Gibco, cat 11965092) supplemented with 10% FBS and 1% PS. HK-2 was cultured in DMEM/F12 (Gibco, Cat 11330032) supplemented with 10% FBS and 1% PS.

##### *Primary cancer cells*

Primary cancer cells were isolated from fresh human ccRCC tissues according to the procedures in previous studies [1]. ccRCC tissue specimens were collected into cold DMEM/F12 medium after nephrectomy until processing within 12 hr. Cancerous tissues were cut into 1 $mm^3$ fragments and digested with 10 mg/mL collagenase type II (Gibco, cat 17101015) in 10 mL HBSS (Gibco, cat 14025092) media, supplemented with 10 μM Y-27632 (Selleckchem, cat S1049) and 1% PS, for about 1 hr at 37 °C, actively vortexed every 15 minutes. After washed 3 times with cold phosphate buffered saline (PBS) (Gibco, cat 10010023), the cell clumps from tumor tissues were then digested with 5ml TrypLE Express (Gibco, cat 12605010), supplemented with 10 μM Y-27632 and 1% PS, for another 5 minutes at 37 °C with constant shaking. Next, primary cancer cells were filtered with 70-μm cell strainer and plated in 10-cm Petri dishes with 20% FBS of culture medium. The Petri dishes were incubated with 1.5% Matrigel in 37 °C incubator for 2 hr in advance. The culture medium for primary cells was consist of DMEM/F12 (Gibco, cat 11330032) supplemented with 10% FBS, 1% GlutaMAX (Gibco, cat 35050061), 1% HEPES (Gibco, cat 15630080), 5 ng/mL FGF2 (PeproTech, cat 100-18B), 5 ng/mL EGF (PeproTech, cat AF-100-15), 5 μg/mL insulin (Sigma, cat I9278), 5 μg/mL transferrin (Sigma, cat T8158), 25 ng/mL hydrocortisone (Apexbio LLC, cat B1951) 10 μM Y-27632 and 1% PS. All experiments were conducted on cells at the first passage.

**Reagents**

LY294002 (Cat S1105), MK2206 (Cat S1078), rapamycin (Cat S1039), and avasimibe (Cat S2187) were purchased from Selleckchem. Cholesteryl oleate (Cat C9253), glyceryl trioleate (Cat T7140), and polybrene (Cat TR-1003-G) were purchased from Sigma-Aldrich.

**Label-free Raman spectromicroscopy**

Label-free Raman spectromicroscopy was performed on unstained frozen tissue slices (~20 μm) and live cells. Compositional analysis of individual LDs and autofluorescent granules was conducted by integration of SRS imaging and confocal Raman spectral analysis on a single platform. SRS imaging was performed on a picosecond SRS microscope. A picosecond pulse laser with 80 MHz repetition rate (Applied Physics & Electronics, picoEmerald™ S) provided the pump and Stokes beams, which were overlapped in space and time. The tunable wavelength of pump beam ranged from 700 nm to 960 nm and the wavelength of Stokes beam was fixed at 1031 nm. Stokes beam was modulated at ~20 MHz by an electronic optic modulator. After combination, two collinear beams were coupled to a two-dimensional scanning galvanometer (Thorlab, GVS012-2D) and then sent to an inverted microscope (Olympus, LX73). A 60x water-immersion objective (Olympus, LUMPlanFL N, NA = 1.0) was employed to focus the light on the sample, and another 60x water-immersion objective (Olympus, LUMPlanFL N, NA = 1.0) was employed to collect the signal. After a short-pass filter (Chroma, ET980SP), the pump beam signal was detected by a 10 mm × 10 mm large-area silicon photodiode (Hamamatsu, S3994-01) detector with 48 DC reversed bias voltage, which was then extracted by a digital lock-in amplifier (Zurich Instruments, HF2LI). The analog output signal was acquired by a data-acquisition card (National Instruments, PCIE-6363) and then sent to the computer for image display on the NI LabVIEW 2018 software.

For SRS imaging of the $CH_2$ vibration (2,850 $cm^{-1}$), pump beam was tuned to 796.8 nm. The power of pump beam at the specimen was maintained at ~20 mW and the power of Stokes beam was kept at ~90mW. Average acquisition time for a 400 × 400 pixels SRS image was ~1.6 second. Large-area SRS images of tissues were recorded by a motorized stage and stitched together up to ~700 µm × 700 µm in size. Simultaneously, backward-detected two-photon fluorescence signal was acquired through a 520/40 nm bandpass filter for the imaging of autofluorescent granules in human tissues. No damage was observed during the imaging process. Images were analyzed by ImageJ 1.53e software.

Spontaneous Raman spectra were acquired by a Raman micro-spectrometer (Andor, Shamrock SR-303i-A) equipped with a cooled charge-coupled device (CCD) detector (Andor, DR316B-LDC-DD). The spectrometer was mounted to the side of the microscope. For confocal measurement, a 50 µm slit was used before the spectrometer. Each Raman spectrum was acquired in 30-40 seconds and ranged from 400 to 3,100 $cm^{-1}$. The laser wavelength was ~707 nm and laser power at the specimen was maintained at ~25 mW. No damage was observed. 6-10 spectra of LDs or autofluorescent granules were obtained in different locations for each specimen. The background of Raman spectra was removed in Origin 2017 software according to the procedures described in the previous study [2].

We quantitatively analyzed LD amount and CE percentage in individual LDs. By using ImageJ “Threshold” function, LDs can be extracted based on the significantly higher signal intensities compared to other cellular compartments. Then, by using ImageJ “Analyze Particles” function, fractions of LD area out of cell area and tissue area were acquired respectively. For tissue specimen of each patient, 6 different locations were averaged for the final value of LD area fraction. For cultured cells, about 20 cells were averaged for the final value of LD amount. For quantification of CE percentage, we then performed SRS imaging and Raman spectral analysis of a series of different molar ratios of TG/CE mixed emulsions ranging from 0:10 to 10:0 as described in previous literatures. The percentage of CE in individual LDs was linearly correlated with the height ratio of the 702 $cm^{-1}$ peak to the 1,442 $cm^{-1}$ peak ($I_{702}/I_{1442}$). Specifically, $I_{702}/I_{1442} = 0.003 \times$ CE percentage (%) (Fig. S2b). CE percentage for each tissue specimen or cell culture was obtained by averaging the CE percentage of LDs in 6-10 cells.

***Hyperspectral SRS microscopy and related quantitative analysis***

For hyperspectral SRS imaging, a dual-output femtosecond (fs) laser source (Insight X3, Spectra-Physics) providing two synchronized beams with a repetition rate of 80 MHz was employed for imaging. The pump beam is tunable from 680 nm to 1300 nm with a pulse width of 120 fs, and the Stokes beam is fixed at 1045 nm with a pulse width of 220 fs. An acoustic-optic (AOM, 1205C, Isomet) modulates the Stokes beam at a frequency of 2.8 MHz. The two beams are collinearly combined through a dichroic mirror (DMSP950, Thorlabs). After the combination, beams are chirped by four 100 mm SF11 glass rods (Newlight) and delivered to a commercial upright laser scanning microscope. Then, a 60X oil immersion objective (NA=1.3, UPlanApo, Olympus) is used to focus two beams on the sample, and an oil immersion condenser (NA=1.4, U-AAC, Olympus) is used to collect the beams from the sample. Two filters (ET980SP, ET795/150 m, Chroma) are used to filter out the Stokes beam, the pump beam is detected by a photodiode (S3994-01, Hamamatsu), and the pump beam loss is extracted by a lock-in amplifier (HF2LI, Zurich Instrument).

For quantification of LD amount, we employed spectral phasor analysis to perform LD detection according to the previous literatures (Anal. Chem. 2014, 86:4115−4119; iScience 2020, 23:100953)[3,4]. We then used the ratiometric approach based on the height ratio of the 2,870 $cm^{-1}$ peak to the 2,850 $cm^{-1}$ peak ($I_{2870}/I_{2850}$) to quantify the CE percentage according to the literature (J. Phys. D: Appl. Phys. 2021, 54:484001)[5]. The measurements of the TG/CE emulsions and establishment of calibration curve were shown in Fig. S5a-S5c. The percentage of CE was linearly correlated with the height ratio $I_{2870}/I_{2850}$. Specifically, $I_{2870}/I_{2850} = 0.0018 \times$ CE percentage (%) + 1.019. The quantification procedure was briefly illustrated in Fig. S5d. For tissue specimen, 6 different locations were averaged for the final value of LD area fraction. For cultured cells, about 20 cells were averaged for the final value of LD amount. CE percentage for each tissue specimen or cell culture was obtained by averaging 6 images.

**Fluorescence Imaging of Dil-LDL/HDL Uptake**

After treated with indicated inhibitors for a given time period, cells were incubated with 20 μg/ml Dil-labeled LDL/HDL (Dil-LDL/HDL) (Yiyuan Biotechnologies, cat YB-0011; Solarbio, cat H8910) for 3 hr at 37 °C in culture medium without FBS and then imaged by two-photon fluorescence microscopy through a 600/60 nm bandpass filter. The Dil-LDL or HDL intensity was quantified with ImageJ and normalized by cell number for lipoprotein uptake capacity.

**Liquid chromatography-mass spectrometry (LC-MS) measurement of lipid extraction**

Lipids were extracted from tissues for LC-MS analysis according the following protocol. The relative levels of CEs were normalized by tissue weight for comparison between normal adjacent and ccRCC tissues. The fold changes of CE levels were further normalized by the normal adjacent tissue group.

The tissues were weighed and homogenized. 1 ml of chloroform/methanol (3:1, v/v) was added and the mixture was ultrasonicated in an ice-water bath for 1 h. Then, 100 μl water was added and mixed. The mixture was centrifuged for 10 minutes at 13200 r/min at 4 ℃. The liquid phase was obtained and then evaporated using a steam of nitrogen. Subsequently, 300 μl of isopropanol/acetonitrile (1:1, v/v) was used to dissolve the residue. After centrifugation, the supernatant was collected and then analyzed by LC-MS.

Separation for relative quantification of CEs was performed by UltiMate™ 3000 Rapid Separation LC (RSLC) system (Thermo Scientific), which carried out using an ACQUITY HSS T3 C18 column (Waters, 1.8 μm, 100 mm × 2.1 mm). The mobile phase was made up of solvent A (0.1% formic acid in acetonitrile/water (6:4, v/v) containing 10 mM ammonium acetate) and solvent B (0.1% formic acid in acetonitrile/isopropanol (9:1, v/v) containing 10 mM ammonium acetate) with a gradient elution (0-2 min, 20-30% B; 2-5 min, 30-45% B; 5-6.5 min, 45-60% B; 6.5-12 min, 60-65% B; 12-14 min, 65-85% B; 14-17.5 min, 85-100 % B; 17.5-18 min, 100-100% B). The flow rate was set at 0.3 ml/min and the column temperature was kept at 50 ℃. The Q Exactive™ hybrid quadrupole Orbitrap mass spectrometer (Thermo Scientific) equipped with a HESI-II probe was operated in the positive electrospray ionization mode under the following conditions: Heated capillary temperature was kept at 320 ℃. Spray voltage was 3700V. Sheath gas pressure was 30 psi. Auxiliary gas was 10 psi. The resolution of full mass scan was 70000 and the mass acquisition range was from *m*/*z* 80-1200.

**DNA sequencing**

Genomic DNA was extracted and purified from primary cells according to standard manufacturer's instruction of DNAzol™ reagent (Thermo Fisher Scientific, cat 10503027). The following three exons of VHL gene were amplified by polymerase chain reaction (PCR) with Q5 High-Fidelity DNA Polymerase ((New England Biolabs (NEB), cat M0491S) and Sanger sequencing was performed to detect VHL mutation by SinoGenoMax (Beijing, China) using standard protocols. The primer sequences were synthesized by Sangon Biotech Co, Ltd (Shanghai, China) and described in the Supplementary Table S11.

**RNA extraction and RNA sequencing**

The TRIzol™ reagent (Thermo Fisher Scientific, cat 15596026) was employed for the RNA extraction from cells according to manufacturer's instruction. Sequence libraries were generated and sequenced by CapitalBio Technology (Beijing, China) and all samples were sequenced on an Illumina HiSeq sequencer (Illumina). For samples with duplicate groups, DESeq2 was used for differential gene expression analysis to get log2(Fold Change) and p-adjust, while those samples that did not contain duplicate groups (treated with avasimibe) were analyzed by using |log2(Fold Change) |>1 (Fold Change = treat/control group), along with intersection between two independent cases of primary cancer cells, as a criterion to determine differential gene.

**Real-time quantitative PCR (qRT-PCR)**

After total RNA extraction from primary cancer cells, reverse transcription was performed using SuperScript III First-Strand Synthesis SuperMix for qRT-PCR kit (Invitrogen, cat 11752-050) according to manufacturer's instruction. Real-time PCR analysis was then conducted using SYBR™ Green PCR Master Mix kit (ABI, cat 4367659). The experiments were performed in duplicate on at least two patients, and each patient was repeated three times. Glyceraldehyde-3-phosphate dehydrogenase (GAPDH) was used as the internal control. The primers were synthesized by Sangon Biotech Co, Ltd (Shanghai, China) and listed in the Table S11.

**RNA interference**

The HIF1α, HIF2α, SREBP-1, SREBP-2, and control shRNA lentiviral particles were purchased from Santa Cruz (Cat sc-35561-V, sc-35316-V, sc-36557-V, sc-36559-V, sc-108080). The VHL (VHL-OE), SREBP-1 (SREBP1-KD) and SREBP-2 (SREBP1-KD) lentiviral particles were designed and synthesized by GeneChem Co, Ltd (Shanghai, China). Lentiviral particles were introduced into cells with 5 μg/mL polybrene and stable transfected primary ccRCC cells were selected by 2 μg/ml puromycin (Selleckchem, cat S7417).

**Filipin and BODIPY staining**

The filipin complex (MedChemExpress, cat 11078-21-0) was employed for free cholesterol staining with a working solution of 0.05 mg/ml in PBS/10% FBS. BODIPY 493/503 (Invitrogen, cat D2191) was used for LD staining with a working solution of 1 μg/mL in PBS. Cells were fixed with 10% formalin solution for 0.5 hour and then incubated with 1.5 mg/mL glycine in PBS for 10 minutes to quench the formalin. Subsequently, cells were stained with 1 mL of filipin working solution for 2 hours and BODIPY working solution for 0.5 hour in the dark. The fluorescence images were immediately obtained with a confocal fluorescence microscopy (Leica) after washed with PBS three times.

**Immunofluorescence staining**

ccRCC primary cells were fixed with 10% formalin solution for 0.5 hour and permeabilized with 0.2% Triton X-100 (Solarbio, cat T8200) for 15 minutes at room temperature. After washed with PBS, cells were blocked with 5% BSA (Beyotime, cat ST023-200g) for 0.5 hour. The cells were then incubated with a primary antibody: anti-pan Cytokeratin (Abcam, cat ab7753)/ ITGA6 (Abcam, Cat ab235905)/ ITGB1 (Abcam, Cat ab30394)/ Caveolin-1 (Cell Signaling Technology, cat 3267S)/ Caveolin-2 (Cell Signaling Technology, cat 8522S) in 5% BSA for 2 hours, followed by incubation in the appropriate secondary antibody: Goat Anti-Mouse IgG Alexa Fluor 647 (Abcam, cat ab150115)/ Donkey Anti-Rabbit IgG Alexa Fluor 647 (Abcam, cat ab150075) in 5% BSA for 1 hour at room temperature. The fluorescence images were immediately examined with a confocal fluorescence microscopy (Leica) after washed with PBS three times.

**Immunoblotting and antibodies**

Cells were lysed by using RIPA buffer (Thermo Fisher Scientific, cat 89900) supplemented with Protease/Phosphate Inhibitor Cocktail (Cell Signaling Technology, cat 5872S). Western blots were performed according to standard procedures and using following antibodies: β-actin (Cat 4970S), VHL (Cat 68547S), HIF1α (Cat 3716S), HIF2α (Cat 59973S), p-S6 (Cat 2211S), and p-AKT (Cat 4060S) were purchased from Cell Signaling Technology; SCARB1 (Cat ab52629), LDL Receptor (Cat ab30532), SREBP-1 (Cat ab3259), SREBP-2 (Cat ab30682), HMGB1 (Cat ab18256), ITGA6 (Cat ab235905), and ITGB1 (Cat ab30394) were purchased from Abcam. Primary antibodies were detected using HRP-conjugated secondary antibodies (anti-rabbit: Cell Signaling Technology, cat 7074S, RRID:AB_2099233; anti-mouse: Cell Signaling Technology, cat 7076S) and followed by SuperSignal West Femto Maximum Sensitivity Substrate (Thermo Fisher Scientific, cat 34095) and exposure to Mini Chemiluminescent Imaging and Analysis System (Sage Creation Science).

**Cell viability assay**

Cells were grown in 96-well plates (5000 cells/well of cell lines and 15,000 cells/well of primary cells) for 1 day and followed by indicated treatment for 3 days. Cell viability was performed with the MTT colorimetric assay (Thermo

Fisher Scientific, cat M6494). IC50 was obtained by fitting the data with the sigmoidal dose response model in Origin 2017 software.

**Cell cycle analysis**

Primary cells were treated with or without 15 μM avasimibe for 2 days, followed by fixation and stained with 50 μg/mL propidium iodide (PI) (Thermo Fisher Scientific, cat P3566) at 37 °C for 30 min. The DNA content was measured by flow cytometer (Peking University Health Science Center). Cell-cycle phases were analyzed by ModFit software.

**Migration/Invasion assay**

Migration and invasion assays were performed in Transwell chambers (Corning, cat 3422). The upper wells of 8-μm-pore-sized membranes for invasion assay were coated with Matrigel (Corning, cat 354234) for 2 hr in advance, while membranes for migration assay did not need coating. Primary cells were treated with or without 15 μM avasimibe for 3 days and $25\times10^4$ cells were seeded in the upper chamber of the transwells with serum-free media for 9 hr at 37 °C, while the culture media in the lower chamber contain 20% FBS and EGF. The upper chamber of the transwells were then fixed and stained with PI. Confocal fluorescence imaging was performed for 7 locations. The number of PI-stained cells were finally counted.

**PDX mouse model**

All animal care and experimental procedures were performed in accordance with the guidelines for care and use of laboratory animals. Patient-derived tumor xenograft (PDX) mouse model of ccRCC was established according to a previously described protocol [6]. After the removal of necrotic tissues, fresh tumor specimens were partitioned into $3 \times 3 \times 3$ mm$^3$ tissue fragments and subcutaneously transplanted into the right flanks of female NOD-SCID mice (6 to 8 weeks, Shanghai Model Organisms Center, Inc). Tumor volume and body weight were monitored weekly. When the stable xenografts were ~1000 mm$^3$, mice were sacrificed and tumors were subsequently implanted into new NOD-SCID mice.

***Anti-tumor efficacy***

Once the xenografts at the third generation steadily grew to ~150 mm$^3$, mice were randomly divided into two groups (10 mice for each group): vehicle (5% DMSO+30% PEG 300+5% Tween 80+ PBS) and avasimibe (15 mg/kg) groups, and administrated via daily intraperitoneal injections. Tumor size and body weight were measured and recorded twice a week. Tumor volume was calculated using the formula: tumor volume = $0.5 \times$ length $\times$ (width)$^2$ mm$^3$.

Relative tumor volume = tumor volume / initial tumor volume. During the experiment study, the maximal tumor size must be less than 20 mm in any dimension. After 24 days of administration, tumor tissues were collected for label-free Raman spectromicroscopy measurements and histological analysis by H&E and TUNEL staining. Pathological examination was performed by experienced pathologists.

***In vivo safety evaluation***

At the end of 24-day treatment, three mice from each group were sacrificed to collect the vital organs (including in liver, lung, heart, spleen, kidney, and adrenal gland) for histological analysis by H&E staining. Blood samples of all mice were collected for hematological and blood biochemical analysis (including in WBC, RBC, PLT, ALT, AST, and CRE).

***Pharmacokinetic study***

Female NOD-SCID mice without tumor (n = 9) were injected intraperitoneally of avasimibe at a dose of 15 mg/kg. These mice were randomly divided into 3 groups (3 mice per group). At 5 minutes, 0.25, 0.5, 1, 2, 4, 6, 12, and 24 hours post-injection, blood samples of each group were collected via the postorbital venous plexus veins in turn. Avasimibe concentration in blood samples was determined using LC-MS (Shimadzu, LC20ADXR), equipped with an AQ-C18 column (2.1×50mm, 5 μm).

***Biodistribution of avasimibe***

After 0.5 hour post intraperitoneal injection with 15 mg/kg avasimibe, tumors, vital organs, and feces in tumor-bearing NOD-SCID mice (n = 3) were harvested. These samples then were homogenized and quantified by LC-MS to measure avasimibe concentration.

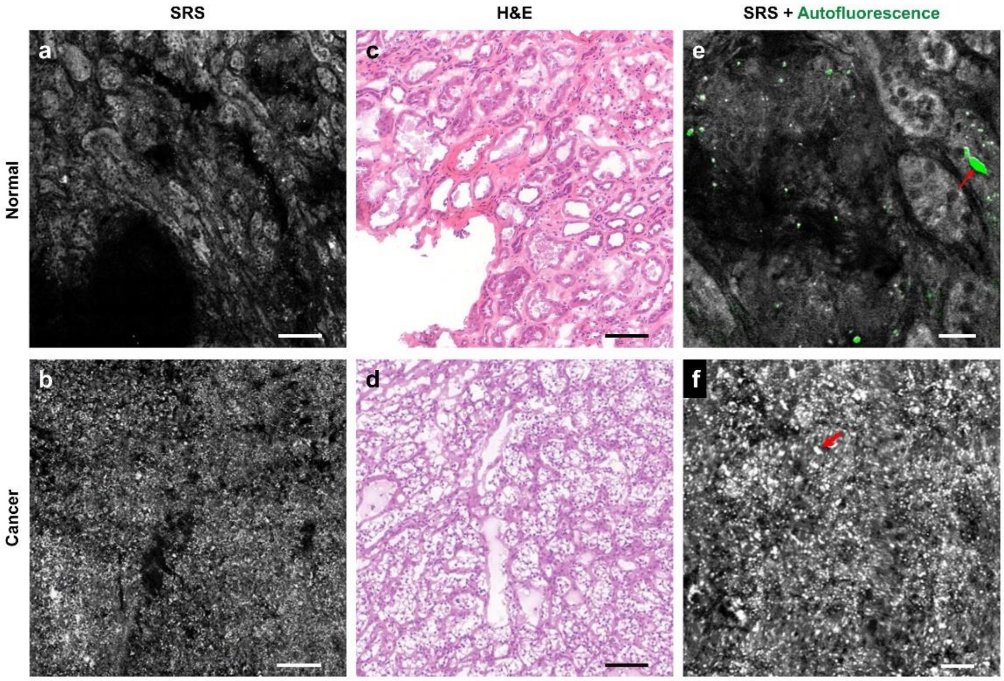


**Figure S1. (related to Fig. 1)**

(**a**, **b**) Representative SRS images of human normal adjacent and ccRCC tissues. Scale bar, 100 µm. (**c**, **d**) H&E images of the adjacent slices shown in **a** and **b**. Scale bar, 100 µm. (**e**, **f**) Magnified SRS and autofluorescence images of the ones shown in **a** and **b**. Autofluorescent granules and LDs are indicated by red arrows. Scale bar, 25 µm.

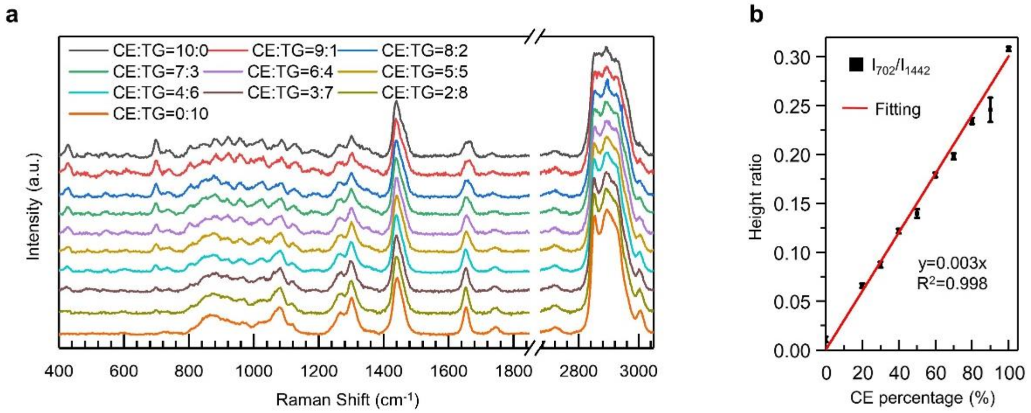


**Figure S2. (related to Fig. 1)**

**(a)** Raman spectra of emulsions mixed with CEs and TGs. Cholesteryl oleate and glyceryl trioleate are mixed in ten different molar ratios, ranging from 10:0 to 0:10. Raman spectral intensity was normalized by the peak at 1,442 $cm^{-1}$. **(b)** Calibration curve for quantification of CE percentage in the emulsions, generated by the ratio of Raman intensity at 702 $cm^{-1}$ to that at 1,442 $cm^{-1}$ ($I_{702}/I_{1442}$). Height ratio = 0.003 × CE percentage (%). Error bars represent SEM.

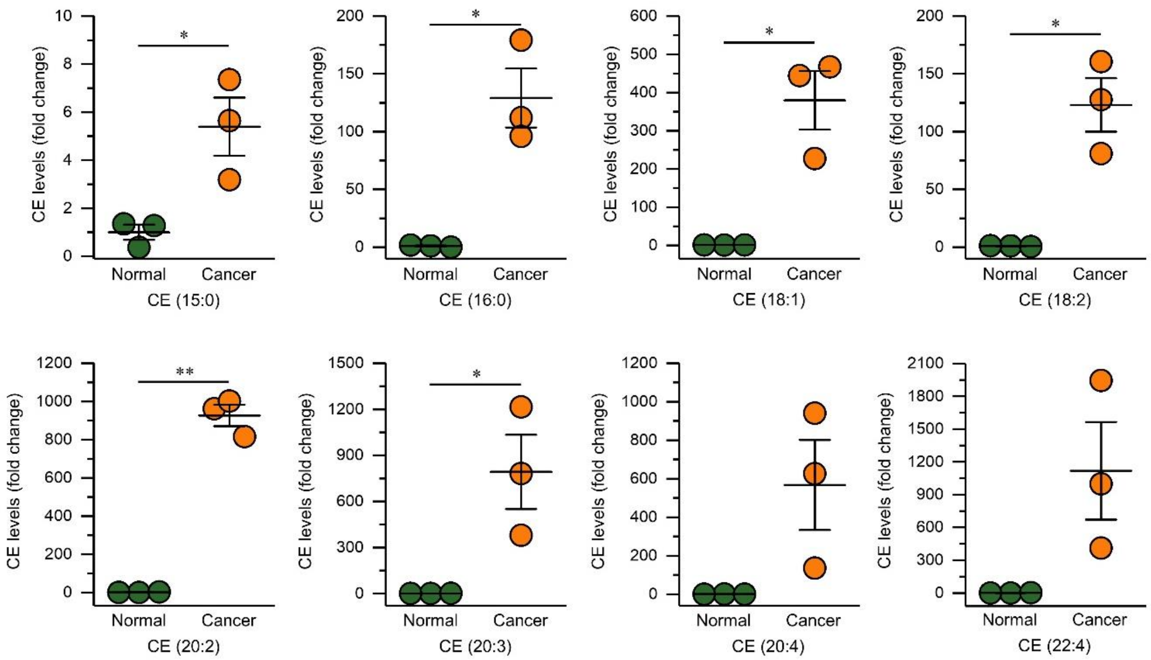


**Figure S3. (related to Fig. 1)**

LC-MS measurement of CEs from lipids extracted from normal adjacent and ccRCC tissues. The fold changes of CE levels were normalized by the normal adjacent tissues. Error bars represent SEM (n = 3). *p < 0.05, **p < 0.005.

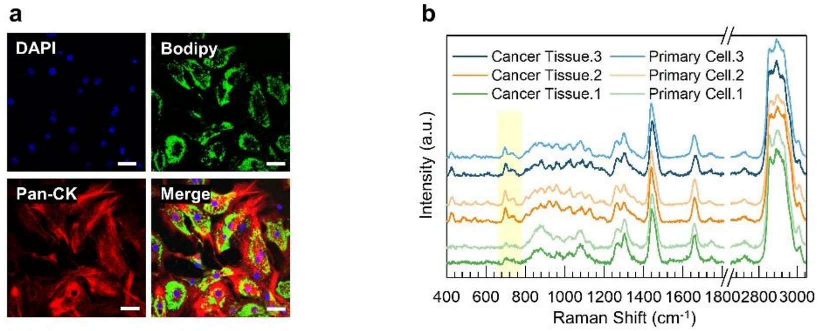


**Figure S4. (related to Fig. 2)**

**(a)** Representative images of DAPI / Bodipy / pan-cytokeratin (pan-CK) staining and the overlapped image of primary cancer cells. Scale bar, 20 µm. **(b)** Representative Raman spectra of LDs in primary cancer cells and their original ccRCC tissues. Spectral intensity was normalized by the peak at 1,442 $cm^{-1}$. Pale yellow area indicates the band of cholesterol rings at 702 $cm^{-1}$.

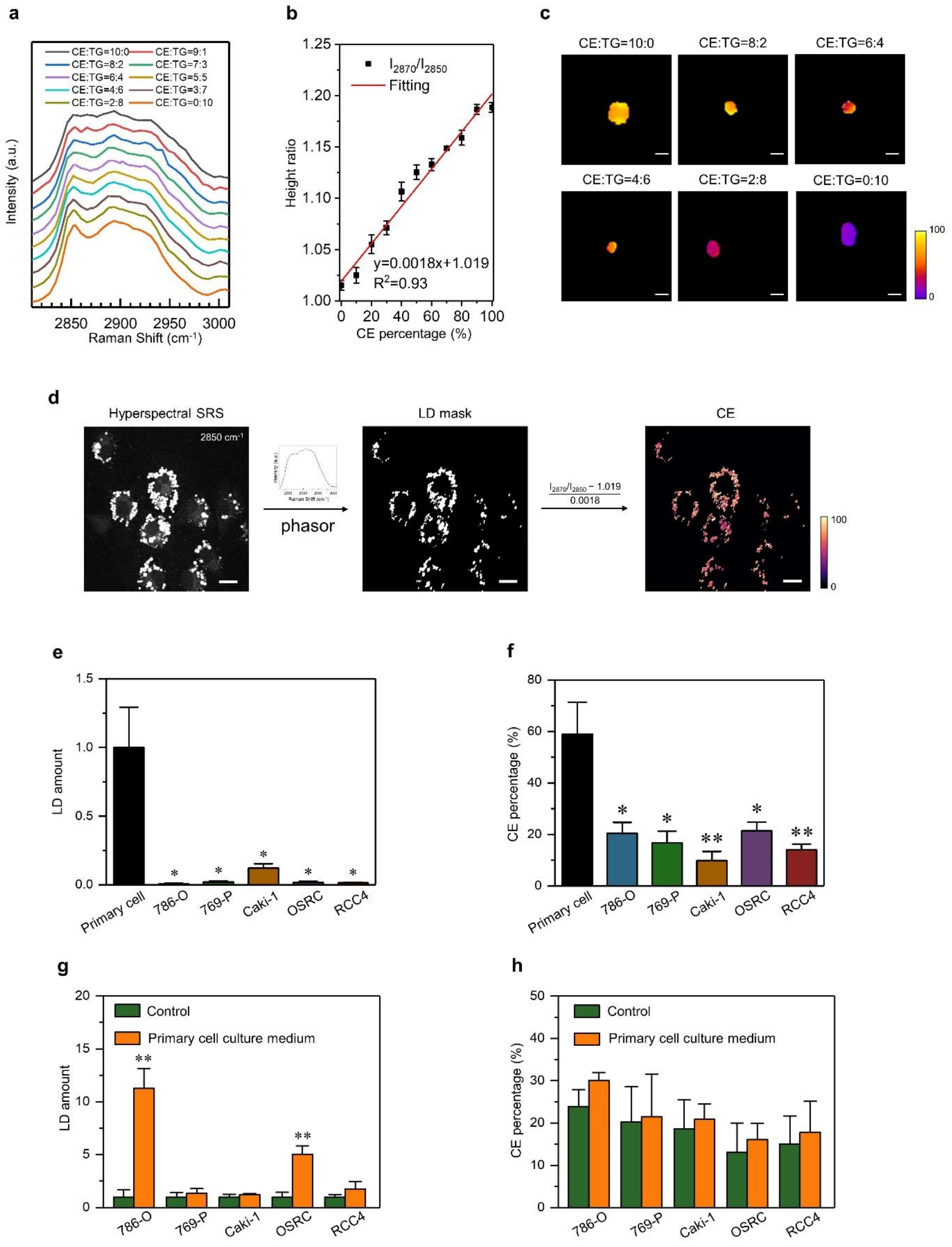


**Figure S5. (related to Fig. 2)**

**(a)** SRS spectra of emulsions mixed with CEs and TGs. Cholesteryl oleate and glyceryl trioleate are mixed in ten different molar ratios, ranging from 10:0 to 0:10. **(b)** Calibration curve for quantification of CE percentage in the emulsions, generated by the height ratio of Raman intensity at 2,870 $cm^{-1}$ to that at 2,850 $cm^{-1}$ ($I_{2870}/I_{2850}$). Height ratio = 0.0018 × CE percentage (%) + 1.019. Error bars represent SEM. **(c)** Quantitative map of CE percentage in six different molar ratios. Scale bar, 5 μm. **(d)** The brief procedure of quantification of LD amount and CE percentage for hyperspectral SRS images. Scale bar, 20 μm. **(e)** Quantitation of LD amount in primary cancer cells and cell lines for hyperspectral SRS imaging, normalized by the LD amount in primary cancer cells. **(f)** Quantitation of CE percentage in primary cancer cells and cell lines for hyperspectral SRS imaging. **(g)** Quantitation of LD amount in the cell lines cultured with primary cell culture medium or regular medium (control), normalized by the LD amount in the control group, by using hyperspectral SRS imaging. **(h)** Quantitation of CE percentage in the cell lines cultured with primary cell culture medium or regular medium (control), by using hyperspectral SRS imaging.

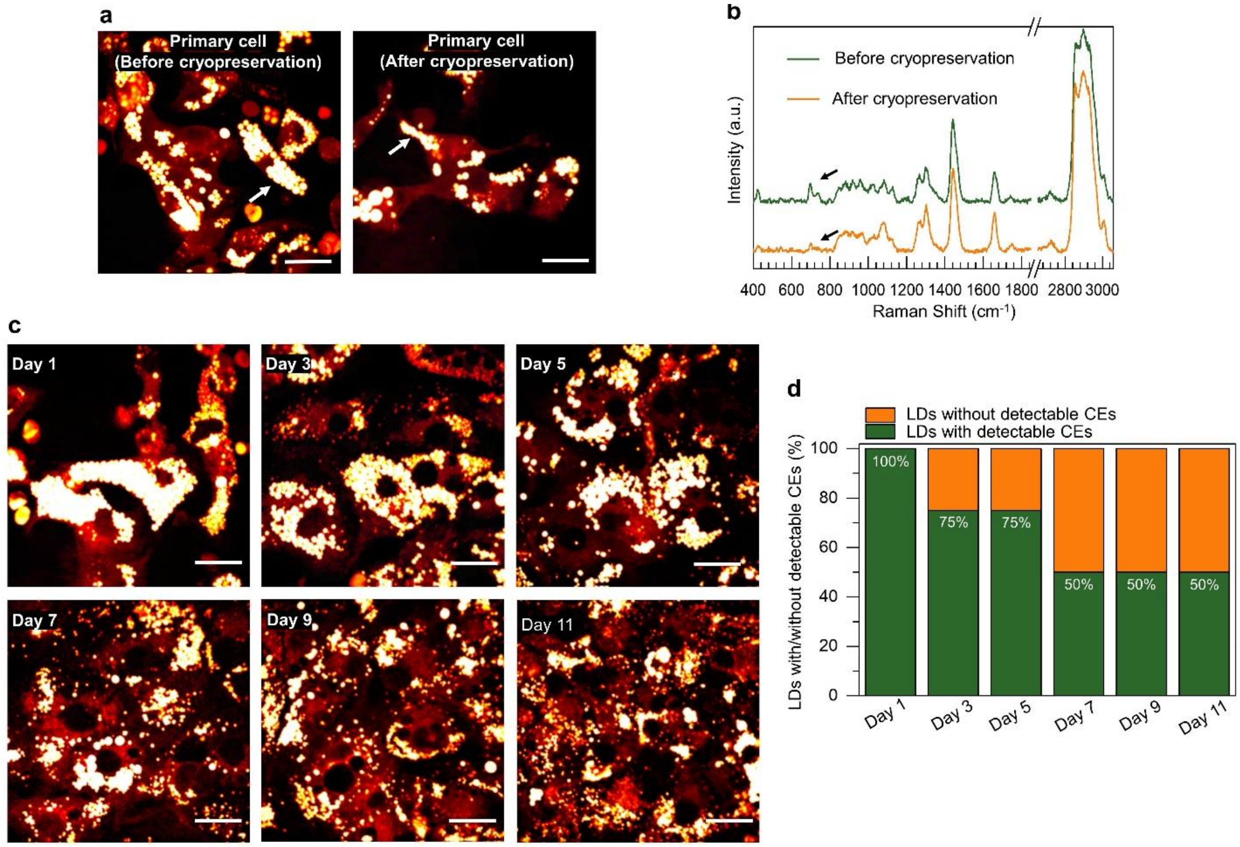


**Figure S6. (related to Fig. 2)**

**(a)** Representative SRS images of primary cancer cells before and after cryopreservation. **(b)** Representative Raman spectra of LDs in primary cancer cells before and after cryopreservation. Spectral intensity was normalized by the peak at 1,442 $cm^{-1}$. The bands of cholesterol rings at 702 $cm^{-1}$ are indicated by black arrows. **(c)** Representative SRS images of primary cancer cells after cultured *in vitro* for 1, 3, 5, 7, 9, 11 days. Scale bar, 20 µm. **(d)** The fraction of LDs with and without detectable CEs in primary cancer cells after cultured *in vitro* for 1, 3, 5, 7, 9, 11 days.

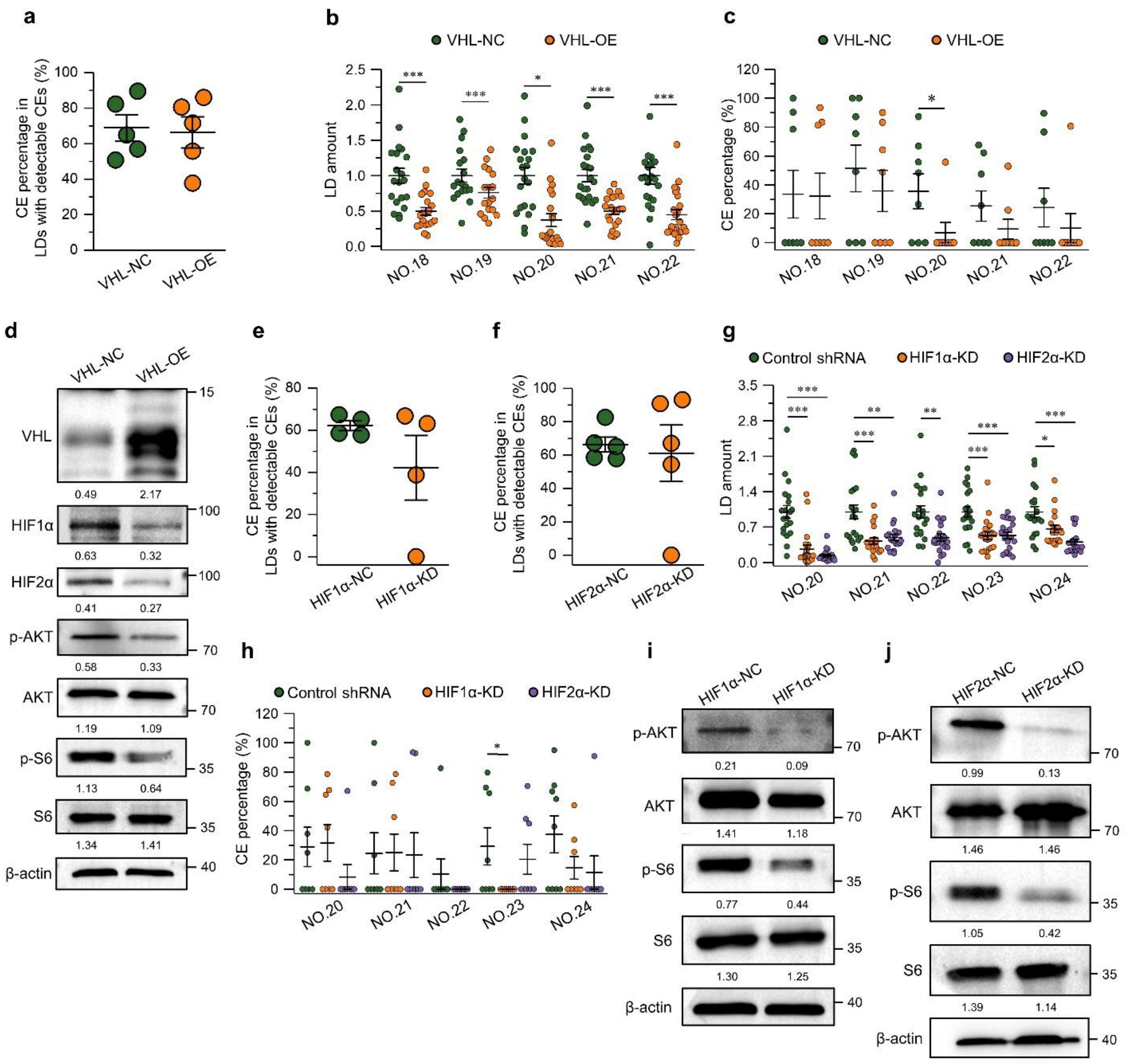

**Figure S7. (related to Fig. 3)**

**(a)** Quantitation of CE percentage of LDs with detectable CEs in primary cancer cells of the VHL-NC and VHL-OE groups. **(b)** Quantification of LD amount in primary cancer cells of the VHL-NC and VHL-OE groups. LD amount was normalized by each VHL-NC group. **(c)** Quantification of CE percentage in primary cancer cells of the VHL-NC and VHL-OE groups. **(d)** Immunoblot of antibodies against VHL, HIF1α, HIF2α, p-AKT, AKT, p-S6, S6, and β-actin in primary cancer cells of the VHL-NC and VHL-OE groups. **(e, f)** Quantitation of CE percentage of LDs with detectable CEs in primary cancer cells of the HIF1α-NC/HIF2α-NC and HIF1α-KD/HIF2α-KD groups. **(g)** Quantitation of LD amount in primary cancer cells of the HIFα-NC, HIF1α-KD, and HIF2α-KD groups. LD amount was normalized by each HIFα-NC group. **(h)** Quantitation of CE percentage in primary cancer cells of the HIFα-NC, HIF1α-KD, and HIF2α-KD groups. Each circle represents average CE percentage of LDs with detectable CEs for one patient in **a**, **e**, and **f**. Each circle represents LD amount for a primary cell in **b** and **g**. Each circle represents CE percentage for a LD in **c** and **h**. Error bars represent SEM. *$p < 0.05$, **$p < 0.005$, ***$p < 0.0005$. **(i, j)** Immunoblot of antibodies against HIF1α/HIF2α, p-AKT, AKT, p-S6, S6, and β-actin in primary cancer cells of the HIF1α-NC/HIF2α-NC and HIF1α-KD/HIF2α-KD groups.

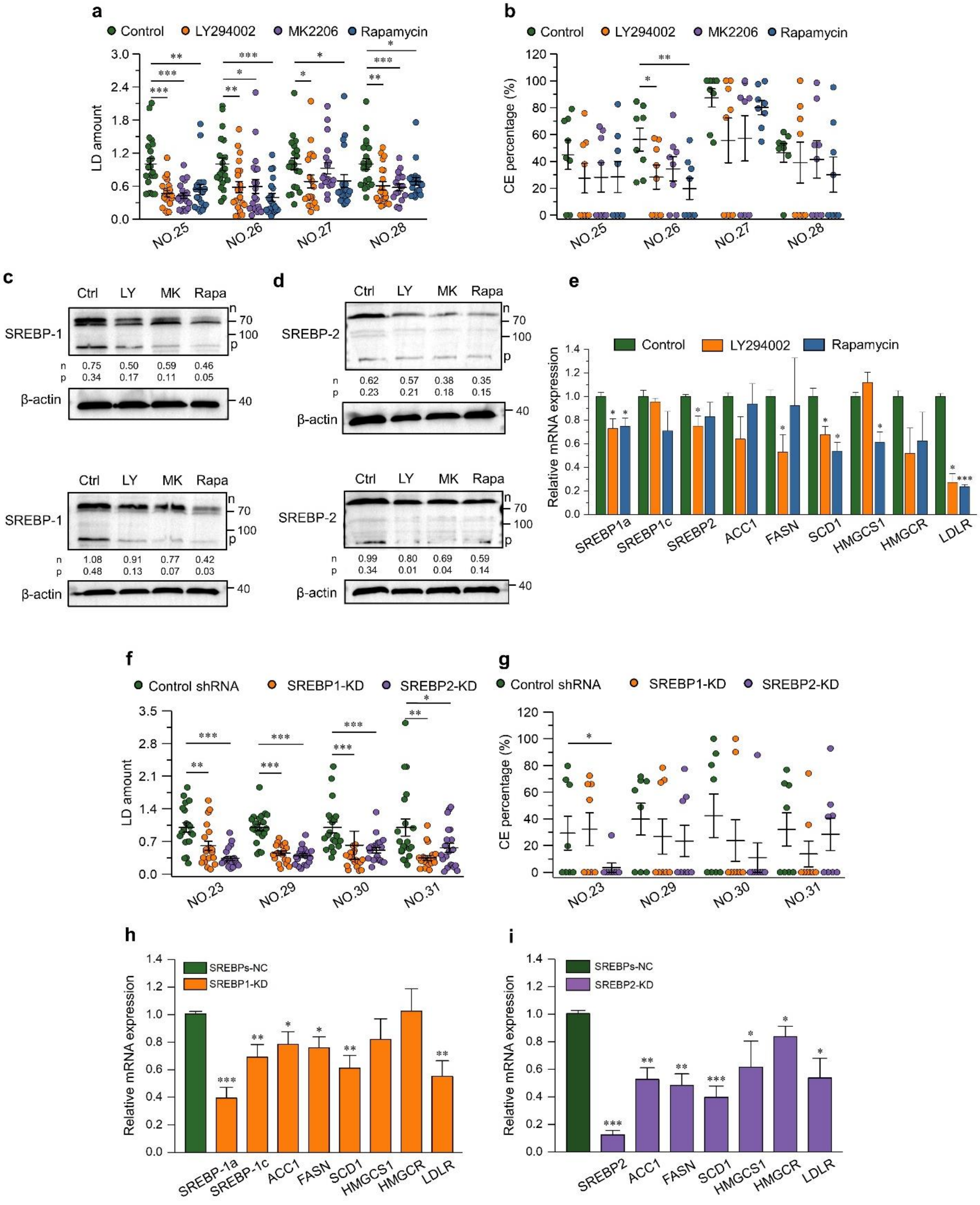


**Figure S8.** **(related to Fig. 4)**

**(a)** Quantification of LD amount in primary cancer cells of control, LY294002, MK2206, and rapamycin groups. LD amount was normalized by each control group. **(b)** Quantification of CE percentage in primary cancer cells of control, LY294002, MK2206, and rapamycin groups. **(c, d)** Immunoblot of antibodies against SREBP-1 (c),

SREBP-2 (d), and β-actin in primary cancer cells treated with DMSO as control (Ctl), LY294002 (50 μM, 3 day) (LY), MK2206 (10 μM, 2 day) (MK), and rapamycin (100 nM, 2 day) (Rapa). p-SREBP, SREBP precursor; n-SREBP, nuclear SREBP. **(e)** Analysis of mRNA expression of SREBP1a, SREBP1c, SREBP2, ACC1, FASN, SCD1, HMGCS1, HMGCR, and LDLR in primary cancer cells of control, LY294002, MK2206, and rapamycin groups via qRT-PCR ($n = 3$). **(f)** Quantification of LD amount in primary cancer cells of the SREBP-NC, SREBP1-KD, and SREBP2-KD groups. LD amount was normalized by each SREBP-NC group. **(g)** Quantification of CE percentage in primary cancer cells of the SREBP-NC, SREBP1-KD, and SREBP2-KD groups. Each circle represents LD amount for a primary cell in **a** and **f**. Each circle represents CE percentage for a LD in **b** and **g**. **(h)** Analysis of mRNA expression of SREBP1a, SREBP1c, ACC1, FASN, SCD1, HMGCS1, HMGCR, and LDLR in primary cancer cells of the SREBP-NC and SREBP1-KD groups via qRT-PCR ($n \geqslant 4$). **(i)** Analysis of mRNA expression of SREBP2, ACC1, FASN, SCD1, HMGCS1, HMGCR, and LDLR in primary cancer cells of the SREBP-NC and SREBP2-KD groups via qRT-PCR ($n \geqslant 4$). Error bars represent SEM. *$p < 0.05$, **$p < 0.005$, ***$p < 0.0005$.

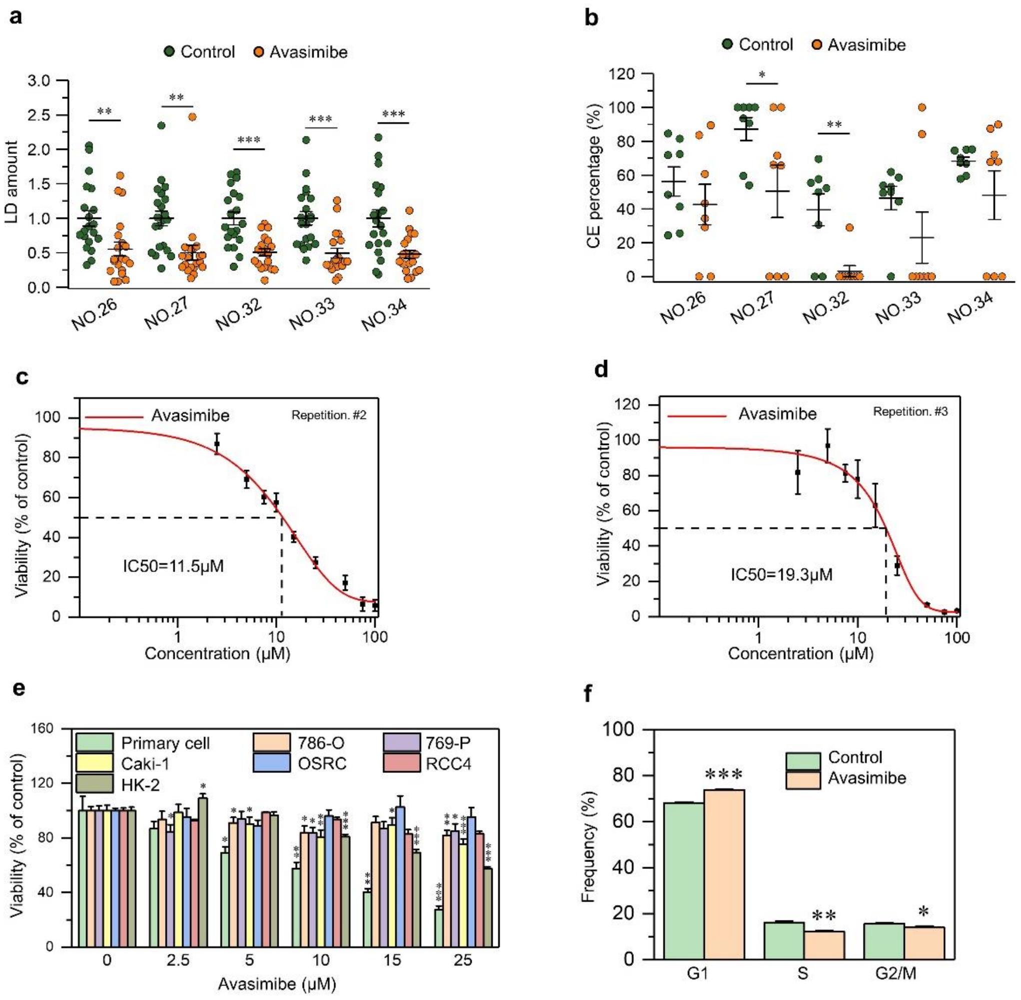


**Figure S9. (related to Fig. 5)**

(**a**) Quantification of LD amount in primary cancer cells of control and avasimibe groups. LD amount was normalized by each control group. Each circle represents LD amount for a primary cell. (**b**) Quantification of CE percentage in primary cancer cells of control and avasimibe groups. Each circle represents CE percentage for a LD. (**c**, **d**) IC50 curve of primary cancer cells upon avasimibe treatment (3 days) in 2 independent patients. (**e**) Viability of primary cancer cells and cell lines (786-O,769-P, Caki-1, OSRC, RCC4, and HK-2) treated with different concentrations of avasimibe (3 days). **(f)** Flow cytometry analysis of cell cycle in primary cancer cells treated with DMSO as control and avasimibe (15 μM, 2 day) (n = 3).

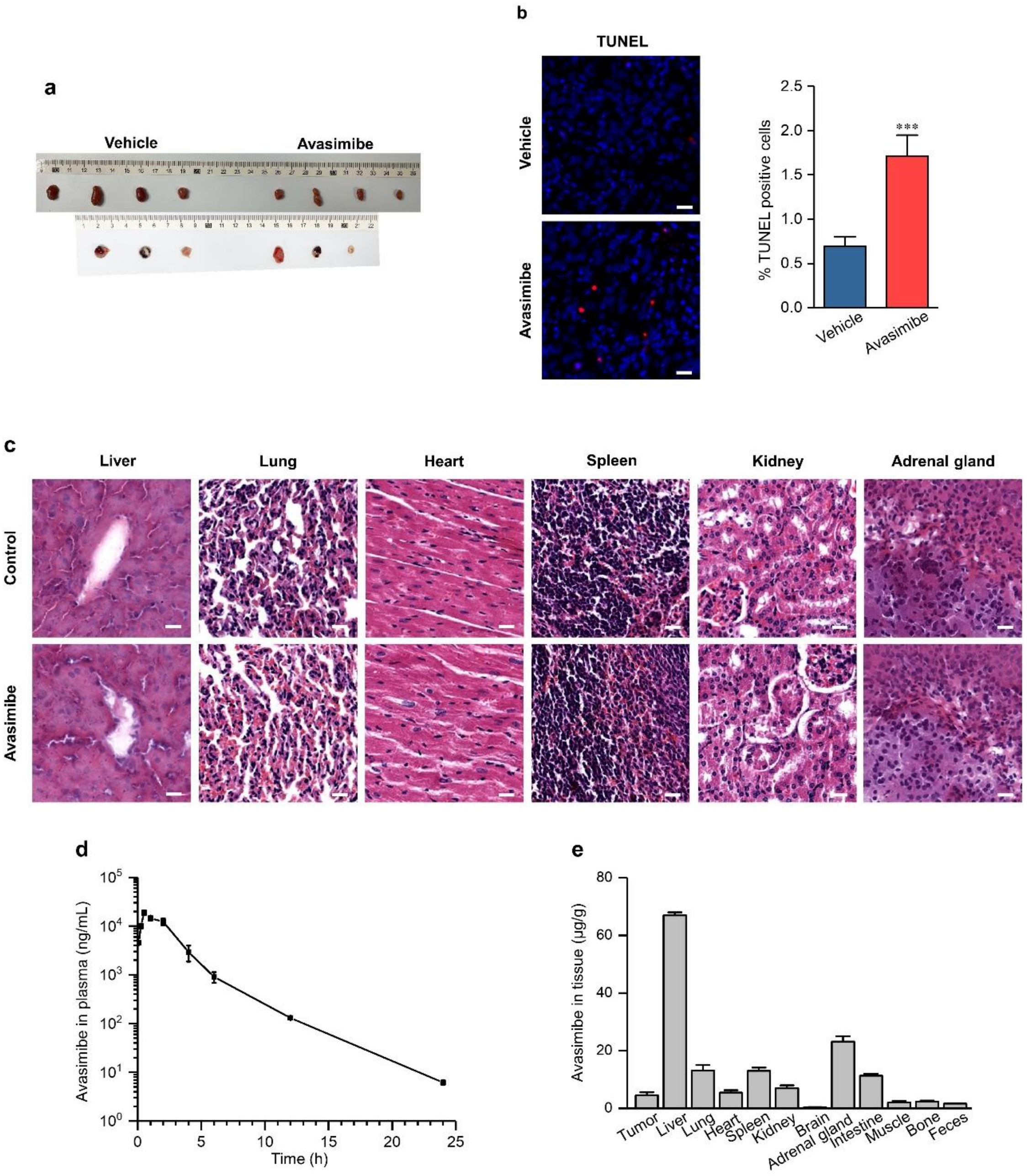


**Figure S10. (related to Fig. 5)**

**(a)** Representative images of tumor tissues harvested from the vehicle and avasimibe-treated PDX mice (n = 7). **(b)** Representative TUNEL staining images (red color: TUNEL staining; blue color: DAPI) and the percentage of TUNEL-positive cells in tumor tissues harvested from vehicle and avasimibe-treated mice (n = 3). **(c)** Representative images of H&E staining of liver, lung, heart, spleen, kidney, and adrenal gland tissues harvested at the end of 24-day treatment of the mice. Scalar bar, 20 µm. **(d)** Mean plasma concentration-time curve of avasimibe at 5 min, 0.25, 0.5, 1, 2, 4, 6, 12, and 24 hours post intraperitoneal injection of avasimibe (n = 3). **(e)** Tissue distribution of avasimibe after 0.5 hour post intraperitoneal injection of avasimibe (n = 3). Error bars represent SEM. *p < 0.05, **p < 0.005, ***p < 0.0005.

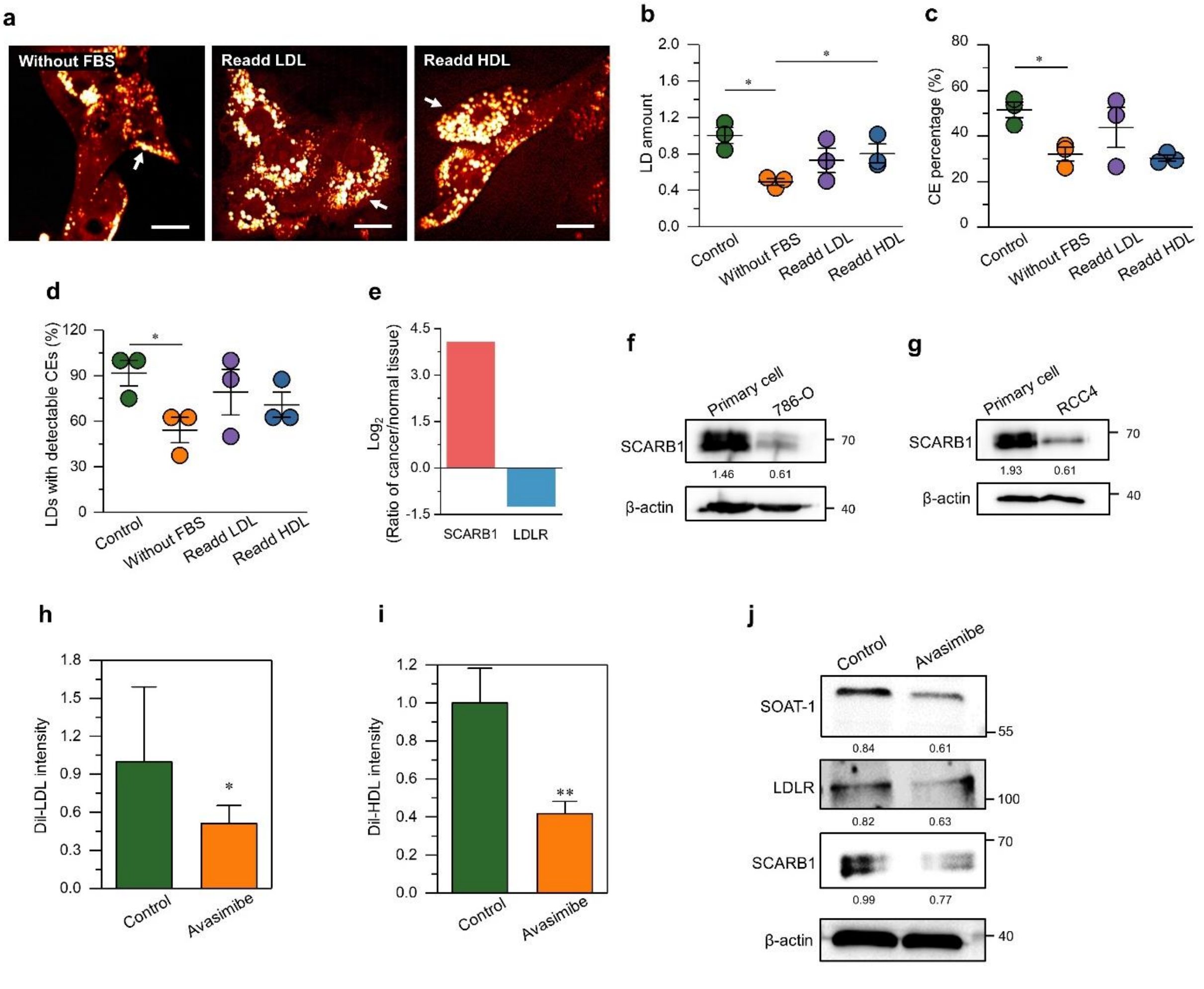


**Figure S11. (related to Discussion)**

**(a)** Representative SRS images of primary cancer cells cultured without FBS (1 day) and then with re-addition of LDL or HDL (45 mg/ml, 1 day). LDs are indicated by white arrows. Scale bar, 20 μm. **(b)** Quantitation of LD amount in primary cancer cells of control, without FBS, and then with re-addition of LDL or HDL. LD amount was normalized by the control group. **(c)** Quantitation of CE percentage in primary cancer cells of control, without FBS, and then with re-addition of LDL or HDL. **(d)** The fraction of LDs with detectable CEs in primary cancer cells of control, without FBS, and then with re-addition of LDL or HDL. **(e)** Significantly differential expression of genes involved in the cholesterol uptake pathway in ccRCC tissues compared to normal tissues based on TCGA dataset ($p$-value < 0.05 and $|\log2(\text{cancer/normal tissues})| > 0.5$). **(f, g)** Immunoblot of antibodies against SCARB1 and β-actin in 786-O (f), RCC4 (g), and primary cancer cells. **(h, i)** Quantitation of DiI-LDL or Dil-HDL uptake in primary cancer cells of control and avasimibe groups (n = 3). DiI-LDL/HDL intensity was normalized by the control group. Error bars represent SEM. *$p < 0.05$, **$p < 0.005$. **(j)** Immunoblot of antibodies against SOAT-1, LDLR, SCARB1, and β-actin in primary cancer cells of control and avasimibe groups.

**Table S1. Presence of autofluorescent granules, area fraction of LDs and CE percentage in normal adjacent and ccRCC tissues shown in Fig. 1(g)**

| | Autofluorescent granule | LD area fraction (%) | LD area fraction (%) Mean ± SEM | CE percentage (%) | CE percentage (%) Mean ± SEM |
|---|---|---|---|---|---|
| **Normal tissues (n=24)** | All positive | Not detectable | | | |
| **ccRCC tissues (n=24)** | All negative | 2.0 | 13.7 ± 2.1 | 91.0 | 66.5 ± 5.7 |
| | | 23.5 | | 97.6 | |
| | | 7.3 | | 46.4 | |
| | | 28.9 | | 46.6 | |
| | | 27.2 | | 75.0 | |
| | | 7.5 | | 73.4 | |
| | | 3.8 | | 76.6 | |
| | | 17.0 | | 81.6 | |
| | | 27.6 | | 57.8 | |
| | | 4.1 | | 96.9 | |
| | | 9.5 | | 23.4 | |
| | | 25.1 | | 86.4 | |
| | | 18.4 | | 72.4 | |
| | | 23.6 | | 75.4 | |
| | | 6.2 | | 92.6 | |
| | | 21.9 | | 60.8 | |
| | | 9.6 | | 7.0 | |
| | | 4.2 | | 86.0 | |
| | | 6.5 | | 80.6 | |
| | | 3.0 | | 74.3 | |
| | | 9.0 | | 30.9 | |
| | | 2.8 | | 72.0 | |
| | | 9.8 | | 0.0 | |
| | | 31.2 | | 92.5 | |

**Table S2. VHL status, LD amount and CE percentage in primary cancer cells (Related to Fig. 3)**

| Patient ID | Mutation type | Exon | Nucleotide changes | Amino acid changes | LD amount (Mean ± SEM) | CE percentage (%) (Mean ± SEM) |
|---|---|---|---|---|---|---|
| ***1*** | Frameshift | 2 | c.405_408delATTT | p.Tyr135FS | 1.1 ± 0.1 | 60.1 ± 3.1 |
| ***2*** | Frameshift | 3 | c.516delinsAAGCCA | p.Pro172FS | 1.0 ± 0.1 | 60.4 ± 5.3 |
| ***3*** | Missense | 1 | c.251T>A | p.Val84Glu | 1.2 ± 0.1 | 39.8 ± 8.1 |
| ***4*** | Frameshift | 2 | c.374_375delAC | p.His125FS | 1.1 ± 0.1 | 66.7 ± 5.5 |
| ***5*** | Frameshift | 2 | c.399_401delTGA | p.Thr133FS | 0.9 ± 0.1 | 72.3 ± 3.4 |
| ***6*** | Frameshift | 1 | c.274_280del7 | p.Asp92FS | 2.1 ± 0.1 | 76.7 ± 3.9 |
| ***7*** | In-Frame | 3 | c.496_504del9 | p.Val166FS | 1.2 ± 0.1 | 76.6 ± 6.1 |
| ***8*** | Frameshift | 1 | c.240_255del16 | p.Ser80FS | 1.6 ± 0.1 | 88.8 ± 1.7 |
| ***9*** | Frameshift | 1 | c.282_283insG | p.Pro95FS | 1.7 ± 0.1 | 82.8 ± 4.0 |
| ***10*** | Frameshift | 1 | c.226_227delTT | p.Phe76FS | 0.7 ± 0.1 | 69.2 ± 4.0 |
| ***11*** | Frameshift | 1 | c.262_269dupTGCCCGTA | p.Trp88FS | 1.6 ± 0.1 | 58.5 ± 4.2 |
| ***12*** | Not detectable | - | - | - | 0.9 ± 0.1 | 0.0 ± 0.0 |
| ***13*** | Not detectable | - | - | - | 1.2 ± 0.1 | 19.5 ± 8.1 |
| ***14*** | Not detectable | - | - | - | 1.7 ± 0.1 | 39.9 ± 14.8 |
| ***15*** | Not detectable | - | - | - | 0.8 ± 0.1 | 0.0 ± 0.0 |
| ***16*** | Not detectable | - | - | - | 0.9 ± 0.1 | 51.3 ± 13.1 |
| ***17*** | Not detectable | - | - | - | 0.6 ± 0.1 | 19.2 ± 13.3 |

*LD amount was normalized by the average LD amount in primary cancer cells of the non-mutated VHL group.

*No mutation was detected by Sanger sequencing.

**Table S3. Treatment information corresponding to Patient ID**

| Patient ID<br>Treatment | 18 | 19 | 20 | 21 | 22 | 23 | 24 | 25 | 26 | 27 | 28 | 29 | 30 | 31 | 32 | 33 | 34 |
|---|---|---|---|---|---|---|---|---|---|---|---|---|---|---|---|---|---|
| **VHL Overexpression** | √ | √ | √ | √ | √ | | | | | | | | | | | | |
| **HIF1α Knockdown** | | | √ | √ | | √ | √ | | | | | | | | | | |
| **HIF2α Knockdown** | | | √ | √ | √ | √ | √ | | | | | | | | | | |
| **LY294002/MK2206/Rapamycin** | | | | | | | | √ | √ | √ | √ | | | | | | |
| **SREBP1/2 Knockdown** | | | | | | √ | | | | | | √ | √ | √ | | | |
| **Avasimibe** | | | | | | | | | √ | √ | | | | | √ | √ | √ |

**Table S4. LD amount and CE percentage in primary cancer cells transfected with negative control (VHL-NC) and VHL overexpression (VHL-OE) (Related to Fig. 3)**

| Patient ID | LD amount (Mean ± SEM) VHL-NC | LD amount (Mean ± SEM) VHL-OE | CE percentage (Mean ± SEM) VHL-NC | CE percentage (Mean ± SEM) VHL-OE |
|---|---|---|---|---|
| ***18*** | 1.1 ± 0.1 | 0.5 ± 0.1 | 33.6% ± 17.7% | 32.3% ± 16.9% |
| ***19*** | 1.2 ± 0.1 | 0.9 ± 0.1 | 51.5% ± 17.2% | 35.8% ± 15.2% |
| ***20*** | 0.7 ± 0.1 | 0.2 ± 0.0 | 35.6% ± 12.9% | 7.0% ± 7.5% |
| ***21*** | 0.9 ± 0.1 | 0.5 ± 0.1 | 25.4% ± 11.3% | 9.4% ± 7.3% |
| ***22*** | 1.1 ± 0.1 | 0.5 ± 0.1 | 24.4% ± 14.3% | 10.1% ± 10.8% |

*LD amount was normalized by the average LD amount in primary cancer cells of the VHL-NC group.

**Table S5. LD amount and CE percentage in primary cancer cells transfected with negative control (HIF1α-NC) and HIF1α shRNA (HIF1α-KD) (Related to Fig. 3)**

| Patient ID | LD amount (Mean ± SEM) HIF1α-NC | LD amount (Mean ± SEM) HIF1α-KD | CE percentage (Mean ± SEM) HIF1α-NC | CE percentage (Mean ± SEM) HIF1α-KD |
|---|---|---|---|---|
| ***20*** | 1.1 ± 0.1 | 0.3 ± 0.1 | 28.9% ± 14.3% | 31.6% ± 13.3% |
| ***21*** | 0.7 ± 0.1 | 0.3 ± 0.0 | 24.5% ± 15.0% | 25.1% ± 13.4% |
| ***23*** | 1.1 ± 0.1 | 0.6 ± 0.1 | 29.3% ± 13.5% | 0.0% ± 0.0% |
| ***24*** | 1.1 ± 0.1 | 0.8 ± 0.1 | 37.4% ± 13.4% | 14.6% ± 8.3% |

*LD amount was normalized by the average LD amount in primary cancer cells of the HIF1α-NC group.

**Table S6. LD amount and CE percentage in primary cancer cells transfected with negative control (HIF2α-NC) and HIF2α shRNA (HIF2α-KD) (Related to Fig. 3)**

| Patient ID | LD amount (Mean ± SEM) HIF2α-NC | LD amount (Mean ± SEM) HIF2α-KD | CE percentage (Mean ± SEM) HIF2α-NC | CE percentage (Mean ± SEM) HIF2α-KD |
|---|---|---|---|---|
| *20* | 1.1 ± 0.1 | 0.2 ± 0.0 | 28.9% ± 14.3% | 8.4% ± 9.0% |
| *21* | 0.7 ± 0.1 | 0.4 ± 0.0 | 24.5% ± 15.0% | 23.3% ± 16.3% |
| *22* | 0.8 ± 0.1 | 0.4 ± 0.1 | 10.3% ± 11.0% | 0.0% ± 0.0% |
| *23* | 1.1 ± 0.1 | 0.6 ± 0.1 | 29.3% ± 13.5% | 20.4% ± 11.0% |
| *24* | 1.2 ± 0.1 | 0.5 ± 0.1 | 37.4% ± 13.4% | 11.4% ± 12.1% |

*LD amount was normalized by the average LD amount in primary cancer cells of the HIF2α-NC group.

**Table S7. LD amount and CE percentage in primary cancer cells treated with DMSO as control, LY294002, MK2206 and rapamycin (Related to Fig. 4)**

| Patient ID | LD amount (Mean ± SEM) Control | LD amount (Mean ± SEM) LY294002 | LD amount (Mean ± SEM) MK2206 | LD amount (Mean ± SEM) Rapamycin | CE percentage (Mean ± SEM) Control | CE percentage (Mean ± SEM) LY294002 | CE percentage (Mean ± SEM) MK2206 | CE percentage (Mean ± SEM) Rapamycin |
|---|---|---|---|---|---|---|---|---|
| *25* | 1.4 ± 0.1 | 0.6 ± 0.1 | 0.6 ± 0.1 | 0.5 ± 0.1 | 44.9% ± 11.7% | 27.5% ± 11.8% | 28.2% ± 11. 8% | 28.4% ± 12.4% |
| *26* | 1.1 ± 0.1 | 0.7 ± 0.1 | 0.7 ± 0.1 | 0.5 ± 0.1 | 56.2% ± 9.2% | 28.3% ± 9.6% | 34.5% ± 9.7% | 19.6% ± 8.5% |
| *27* | 0.6 ± 0.1 | 0.4 ± 0.1 | 0.5 ± 0.1 | 0.4 ± 0.1 | 87.2% ± 7.3% | 55.6% ± 17.8% | 57.2% ± 18.0% | 80.1% ± 5.8% |
| *28* | 0.9 ± 0.1 | 0.5 ± 0.1 | 0.5 ± 0.1 | 0.6 ± 0.1 | 46.4% ± 7.4% | 39.1% ± 16.3% | 41.4% ± 14.8% | 30.1% ± 14.0% |

*LD amount was normalized by the average LD amount in primary cancer cells of the control group.

**Table S8. LD amount and CE percentage in primary cancer cells transfected with negative control (SREBP-NC), SREBP1 shRNA (SREBP1-KD) and SREBP2 shRNA (SREBP2-KD) (Related to Fig. 4)**

| Patient ID | LD amount (Mean ± SEM) SREBP-NC | LD amount (Mean ± SEM) SREBP1-KD | LD amount (Mean ± SEM) SREBP2-KD | CE percentage (Mean ± SEM) SREBP-NC | CE percentage (Mean ± SEM) SREBP1-KD | CE percentage (Mean ± SEM) SREBP2-KD |
|---|---|---|---|---|---|---|
| ***23*** | 0.8 ± 0.1 | 0.5 ± 0.1 | 0.3 ± 0.0 | 29.3% ± 13.5% | 32.3% ± 13.2% | 3.5% ± 3.7% |
| ***29*** | 1.4 ± 0.1 | 0.6 ± 0.1 | 0.5 ± 0.1 | 39.8% ± 12.7% | 26.8% ± 14.0% | 23.4% ± 12.5% |
| ***30*** | 1.3 ± 0.2 | 0.4 ± 0.1 | 0.7 ± 0.1 | 42.4% ± 17.4% | 23.8% ± 16.7% | 11.0% ± 11.7% |
| ***31*** | 0.5 ± 0.1 | 0.2 ± 0.0 | 0.3 ± 0.1 | 32.1% ± 13.3% | 13.7% ± 10.4% | 28.4% ± 13.0% |

*LD amount was normalized by the average LD amount in primary cancer cells of the SREBP-NC group.

**Table S9. LD amount and CE percentage in primary cancer cells upon avasimibe treatment (Related to Fig. 5)**

| Patient ID | LD amount (Mean ± SEM) Control | LD amount (Mean ± SEM) Avasimibe | CE percentage (Mean ± SEM) Control | CE percentage (Mean ± SEM) Avasimibe |
|---|---|---|---|---|
| ***26*** | 1.1 ± 0.1 | 0.6 ± 0.1 | 56.2% ± 9.2% | 42.6% ± 12.8% |
| ***27*** | 0.5 ± 0.1 | 0.3 ± 0.1 | 87.2% ± 7.3% | 50.4% ± 16.6% |
| ***32*** | 1.4 ± 0.1 | 0.7 ± 0.1 | 39.5% ± 10.1% | 3.2% ± 3.4% |
| ***33*** | 0.9 ± 0.1 | 0.4 ± 0.1 | 46.4% ± 7.4% | 23.0% ± 16.2% |
| ***34*** | 1.1 ± 0.1 | 0.5 ± 0.1 | 68.3% ± 2.6% | 48.1% ± 15.4% |

*LD amount was normalized by the average LD amount in primary cancer cells of the control group.

**Table S10. Mean values of the pharmacokinetic parameters of avasimibe (Related to Fig. 6)**

| Pharmacokinetic parameters | |
|---|---|
| $C_{max}$ (μg/mL) | 18.8 |
| $T_{max}$ (h) | 0.5 |
| $t_{1/2}$ (h) | 2.53 |
| $MRT_{inf}$ (h) | 2.26 |
| CL/F (mL/min/kg) | 5.02 |
| Vz/F (L/kg) | 1.10 |
| $AUC_{0\text{-}t}$ (μg*h/mL) | 49.76 |
| $AUC_{0\text{-}inf}$ (μg*h/mL) | 49.79 |

*$C_{max}$, maximum plasma concentration; $T_{max}$, time to peak concentration; $t_{1/2}$, terminal half-life; MRT, mean residence time; CL/F, apparent clearance; Vz/F, apparent volume of distribution; AUC, area under the plasma concentration-time curve. n=3.

**Table S11. Primer sequences used in the study are shown as follows**

| Target gene | Forward primer | Reverse primer |
|---|---|---|
| VHL-Exon 1 | 5'-GGTGGTCTGGATCGCGGA-3' | 5'-GGCTTCAGACCGTGCTATCG-3' |
| VHL-Exon 2 | 5'-GTGGCTCTTTAACAACCTTTGC-3' | 5'-CCTGTACTTACCACAACAACCTTATC-3' |
| VHL-Exon 3 | 5'-AGTCTGTCACTGAGGATTTG-3' | 5'-CTGAGATGAAACAGTGTAAG-3' |
| SREBP-1a | 5′-TCAGCGAGGCGGCTTTGGAGCAG-3′ | 5′-CATGTCTTCGATGTCGGTCAG-3′ |
| SREBP-1c | 5′-GGAGGGGTAGGGCCAACGGCCT-3′ | 5′-CATGTCTTCGAAAGTGCAATCC-3′ |
| SREBP2 | 5′-CTCCATTGACTCTGAGCCAGGA-3′ | 5′-GAATCCGTGAGCGGTCTACCAT-3′ |
| ACC1 | 5′-TTCACTCCACCTTGTCAGCGGA-3′ | 5′-GTCAGAGAAGCAGCCCATCACT-3′ |
| FASN | 5′-TTCTACGGCTCCACGCTCTTCC-3′ | 5′-GAAGAGTCTTCGTCAGCCAGGA-3′ |
| SCD1 | 5′-CCTGGTTTCACTTGGAGCTGTG-3′ | 5’-TGTGGTGAAGTTGATGTGCCAGC-3’ |

| | | |
|---|---|---|
| HMGCS1 | 5′-AAGTCACACAAGATGCTACACCG-3′ | 5′-TCAGCGAAGACATCTGGTGCCA-3′ |
| HMGCR | 5′-GACGTGAACCTATGCTGGTCAG-3′ | 5′-GGTATCTGTTTCAGCCACTAAGG-3′ |
| LDLR | 5′-ACGGCGTCTCTTCCTATGACA-3′ | 5′-CCCTTGGTATCCGCAACAGA-3′ |
| GAPDH | 5′-GTCTCCTCTGACTTCAACAGCG-3′ | 5′-ACCACCCTGTTGCTGTAGCCAA-3′ |
